# Enhanced mitochondrial G-quadruplex formation impedes replication fork progression leading to mtDNA loss in human cells

**DOI:** 10.1101/2022.06.08.495283

**Authors:** Mara Doimo, Sanna Abrahamsson, Valentin L’Hôte, Mama Ndi, Rabindra Nath Das, Koit Aasumets, Andreas Berner, Steffi Goffart, Jaakko L.O. Pohjoismäki, Marcela Dávila López, Erik Chorell, Sjoerd Wanrooij

**Author notes:** To whom correspondence should be addressed: Sjoerd Wanrooij, Department of Medical Biochemistry and Biophysics, Umeå University, 90187 Umeå, Sweden.; correspondence may also be addressed to: Mara Doimo, Department of Medical Biochemistry and Biophysics, Umeå University, 90187 Umeå, Sweden.

## Abstract

Mitochondrial DNA (mtDNA) replication stalling is considered an initial step in the formation of mtDNA deletions that associate with genetic inherited disorders and aging. However, the molecular details of how stalled replication forks lead to mtDNA deletions accumulation are still unclear. Mitochondrial DNA deletion breakpoints preferentially occur at sequence motifs predicted to form G-quadruplexes (G4s), four-stranded nucleic acid structures that can fold in guanine-rich regions. Whether mtDNA G4s form *in vivo* and their potential implication for mtDNA instability is still under debate.

In here, we developed new tools to map G4s in the mtDNA of living cells. We engineered a G4-binding protein targeted to the mitochondrial matrix of a human cell line and established the mtG4-ChIP method, enabling the determination of mtDNA G4s under different cellular conditions. Our results are indicative of transient mtDNA G4 formation in human cells. We demonstrated that mtDNA-specific replication stalling increases formation of G4s, particularly in the major arc. Moreover, elevated levels of G4 block the progression of the mtDNA replication fork and cause mtDNA loss. We conclude that stalling of the mtDNA replisome enhances mtDNA G4 occurrence, and that G4s not resolved in a timely manner can have a negative impact on mtDNA integrity.

## Introduction

Mitochondria are key organelles as they are responsible for supplying the proper form of energy necessary for the cell to exert all its functions. Conversely to other cellular compartments, they possess their own circular, multi-copy DNA (mtDNA). The genes encoded by this small genome are essential for the function of the mitochondria (1). Qualitative (deletions) and/or quantitative (depletion) loss of the genetic information harboured in the mtDNA is detrimental for cellular homeostasis and, in human, correlates with the onset of several pathological conditions collectively referred to as mtDNA maintenance defects (MDMD) (2).

One of the two strands of the mtDNA, the so-called “heavy strand” (H-strand), is guanine-rich and harbours several sequences with the potential to form G-quadruplexes (G4s). G4s are non-canonical DNA secondary structures formed in nucleic acid sequences rich in guanines. In the human nuclear genome (nDNA), there are at least 700.000 sequences with the propensity to form G4s (3). G4s forming in nDNA are involved in the regulation of several cellular processes, such as replication, transcription and telomere maintenance (4). In mitochondria, G4s play a role in regulation of gene expression (5–7). In contrast, the *in vivo* distribution of G4s in the mtDNA and their implications in the process of mtDNA replication are still unknown. Nonetheless, *in silico* data showed that putative G4-forming sequences associate with pathogenic mtDNA deletion breakpoints (8, 9).

However, the precise role of G4s in the process of mtDNA deletions formation has remained elusive, but work from the nDNA replication field has demonstrated that G4s may impede the progression of the nDNA replication machinery (10). Biochemical studies indicate that the formation of these structures might also interfere with mtDNA replication (9, 11).

The study of G4s dynamics is limited due to the lack of tools to detect these structures in the cellular context. Recently, several groups developed synthetic antibodies or chemical compounds with fluorescent properties that specifically bind and recognize G4s. The first approach, employing synthetic antibodies, allows G4s detection in cultured cells using conventional immuno-techniques (12). Although G4s-specific antibodies proved to be useful to demonstrate the *in vivo* relevance of G4s, this method is subjected to specific manipulation (e.g. fixation and permeabilization) that can alter the G4 formation and therefore the antibody recognition (13). In addition, this approach does not allow to specifically recognize G4s on the mtDNA without detecting nDNA G4s. In contrast, the second approach, employing chemical compounds, can be used in living cells, but often requires the use of non-conventional microscope techniques (e.g. 2-photon microscopy (13)) and the compounds are generally strictly localized to the nucleus (14). Recently, some groups developed fluorescent compounds specific for mtDNA that allow to follow the dynamic of G4s in living cells. However, these compounds do not permit to identify the specific localization of the G4s in the mtDNA sequence (15–17).

To overcome these limitations, we established a cell model in which the G4-binding synthetic antibody BG4 (12) is specifically localized in the mitochondria and developed a chromatin immunoprecipitation approach to map specifically mtDNA G4s, called mtG4-ChIP. This method allowed to map the regions in the mtDNA that accumulate G4s under physiological conditions. We then apply the mtG4-ChIP protocol to study the interplay between mtDNA G4s formation and the loss of mtDNA integrity.

## MATERIALS AND METHODS

### Cloning

The *MTSBG4* construct was synthesized by two-step overlap extension PCR. The *BG4* sequence was amplified by PCR from the pSANG10-3F-BG4 plasmid (Addgene plasmid # 55756(12)) while the mitochondrial targeting sequence (MTS) of TFAM (aa 1-50) was amplified from the pcDNA5-FRT-TO-MTS-PrimPol construct (18). The reaction for the annealing PCR was performed with 0.5 μL of a two-fold dilution of each reaction mixture from first-step amplifications. All amplifications were performed with Phusion polymerase (NEB) according to the manufacturer’s instructions. The thermal conditions were as follows: 1 min at 98 °C, followed by 20 cycles of 10 sec at 98 °C, 15 sec at 55.8 °C, and 30 s at 72 °C, then 7 min at 72 °C. Oligonucleotide used for cloning are listed in Sup. Table 4. The PCR product was then inserted into the pcDNA5-FRT-TO construct (ThermoFisher Scientific) using *Hind*III and *EcoR*V restriction sites following standard procedures.

### Cell culture and compounds preparation

Flp-In T-REx 293 (ThermoFisher Scientific) were maintained in DMEM high glucose + GlutaMAX (Life Technologies) supplemented with 10 % Fetal Bovine Serum (Sigma-Aldrich) and 1 mM pyruvate in standard incubation conditions at 37 °C and 7 % CO2. Upon integration of the MTS-BG4 gene, 50 μg/mL uridine was added to the media. 2’,3’-dideoxycytidine (ddC, Abcam) was freshly dissolved in distilled water to 100 mM prior to cell treatment. RHPS4 (TOCRIS) was dissolved to 10 mM in distilled water and stored at -20 °C in small aliquots.

### Generation of MTS-BG4 cell lines (ρ+ and ρ0)

The generation of the HEK 293 Flp-In T-REx 293 MTS-BG4 (mitoBG4) cell line was performed as previously described (19). Briefly, pcDNA5-FRT-TO MTS-BG4 was co-transfected with the pOG44 plasmid (ThermoFisher Scientific), containing the Flp recombinase, in a 1:10 ratio in a 80 % confluent 10 cm plate using TurboFect Transfection Reagent. The day after transfection, cells were plated in five plates at differential dilutions and were cultured in the presence of 200 μg/mL hygromycin B for several days until isolated colonies were visible. Hygromycin B resistant cells were pooled and kept in growth medium containing 100 μg/mL hygromycin B. To allow the expression of MTS-BG4, doxycycline was added to the medium at the concentrations indicated for each experiment.

The mitoBG4 ρ0 cell line was generated by keeping the cells in regular high glucose medium supplemented with 150 ng/mL Ethidium Bromide for 45 days. Depletion of mtDNA pools was measured at every cell passage by Multiplex PCR as described below. After complete depletion of mtDNA, cells were maintained on regular high glucose medium supplemented with pyruvate and uridine. Medium was changed every second day to avoid acidification. Before each experiment, cells were checked for complete depletion of mtDNA.

### Protein extraction and immunoblotting

Cells were solubilized for 30 min on ice in RIPA buffer (150 mM NaCl, 1 % NP-40, 0.1 % SDS, 0.5 % sodium deoxycholate, and 50 mM Tris-HCl (pH 8.0)). After high-speed centrifugation, the supernatant was collected for further analysis. Buffers for protein extraction were supplemented with 1×EDTA-free Halt protease inhibitor cocktail (ThermoFisher Scientific). Protein amounts were quantified using a BCA protein assay kit (ThermoScientific). Equal amounts (15 μg) of proteins dissolved in 1x Laemmli Sample Buffer (Biorad) with 2.5 % β-mercaptoethanol were separated on 4−20 % SDS-TGX (Biorad) gels and transferred to 0.45 μM nitrocellulose membranes (GE Healthcare Life Sciences) using a Mini-Protean electrophoresis system (Bio-Rad). Membranes were blocked in 5 % nonfat milk for 2 h. Primary antibodies were incubated overnight at 4 °C, and horseradish peroxidase-conjugated-secondary antibodies were incubated 1 h at room temperature. The antibodies used and their dilutions are listed in Sup. Table 2. All washes and incubations were performed in Tris-buffered saline with Tween-20. Chemiluminescent detection was performed using ECL Western blot substrates (ThermoScientific) and a ChemiDoc Touch Imaging System (Bio-Rad).

For non-reducing SDS-PAGE, samples were extracted in NP-40 lysis buffer (150 mM NaCl, 1 % NP-40, 10 % glycerol and 50 mM Tris-HCl (pH 8.0)) supplemented with 1× EDTA-free Halt protease inhibitor cocktail (ThermoFisher Scientific), 1 mM EGTA (pH 7,7), 5 mM ZnCl, 5 mM NaF and 1 mM Na3OV4. Native samples were dissolved in non-reducing SDS Sample Buffer (10 % SDS, 30 % glycerol, 250 mM Tris HCl (pH6,8)) prior to SDS-PAGE separation and immunoblotting as described above. Recombinant BG4 (recBG4) (Millipore Merck) was added as control.

### Cell fractionation and proteinase K accessibility assay

Cells were seeded on 15-cm plates in order to reach 80 % confluency on the day of fractionation, and treated for 24 h with 10 ng/mL doxycycline. Cell fractionation and proteinase K accessibility assay were performed as previously described (18). Samples were analyzed by immunoblot as described above. Antibodies used are described in Sup. Table 4.

### Immunohistochemistry (IHC) and co-localization analysis

IHC was performed using a protocol modified from Jamroskovic et al, 2020 (20). Briefly, 60 000 cells were seeded on 13 mm glass coverslips the day before treatment and induced with 100 ng/mL doxycycline for 24 h. Cells were fixed in 2 % paraformaldehyde and permeabilized in 0.1 % Triton X-100 at room temperature. Cells were blocked with 10 % goat serum followed by incubation with antibodies diluted in 5 % goat serum. For primary antibodies, 0.1 % TWEEN was added. Each incubation was performed for 1 h at 37 °C in a humidified chamber. The antibodies used and their dilutions are described in Sup. Table 2. All washes and incubations were performed in 1× PBS buffer. Coverslips were mounted on glass slides with DAKO mounting medium (Agilent Technologies) and stored at 4 °C.

Slides were imaged with Leica SP8 FALCON Confocal equipped with HC PL APO 63x/1.40 OIL CS2 objective. Fluorophores were excited sequentially with white light laser (WLL) and recorded with HyD detectors (Alexa Fluor 488: Ex: laser line 499, Em: 510-593nm; Alexa Fluor 594: Ex: laser line 598 Em: 610-782nm). Multiple z-stacks were collected for each field. A zoom of 3 was applied. Settings were optimized for co-localization analysis (21). Mander’s correlation coefficient was measured with ImageJ (22) using the JaCoP plugin (21). Around 50 cells were analyzed for each experiment.

### Long term mitoBG4 expression

Cells were seeded in 6 wells-plate at 200 000 cells/well and treatment with 10 ng/mL doxycycline was started the day after seeding. Cells were split 1:10 every third day and maintained in doxycycline. The remaining cells were used for immunoblot and copy number analysis as described in the respective sections.

### MtDNA copy number

MtDNA copy number was measured by multiplex PCR (23). Cells from the indicated treatment were collected by trypsinization and total genomic DNA was extracted using the PureLink Genomic DNA Minikit (Thermo Fisher Scientific). Each DNA sample was diluted to 1 ng/μL and 2 μL were used for amplification using PrimeTime Gene expression Master Mix (IDT) with LightCycler96 (Roche) instrument. Oligonucleotides for nuclear and mitochondrial DNA amplification and fluorescent labelled probes for signal detection are indicated in Sup. Table 1. Each sample was run in triplicate. Mean Ct values were used to calculate the mtDNA copy number relative to untreated samples using the ΔΔCt method.

### Two-Dimensional AGE (2D-AGE) and Southern blotting

2D-AGE and subsequent Southern blot were performed as previously described (18). HincII (ThermoScientific) was used for mtDNA fragmentation. A PCR probe spanning nucleotides 35-611 (fcorresponding to the mtDNA non-coding region) random-prime labelled with [α-^32^P]dCTP was used for nucleic acid hybridization.

For the analysis of sheared DNA samples for ChIP protocol, reverse cross-linked DNA was precipitated and purify using standard phenol-chloroform extraction. 2,5 μg of non-sheared and sheared DNA were separated over a 1 % agarose gel in 1x TBE buffer. Southern blotting was performed by using standard procedures. An oligonucleotide mapping to the ND5 region of the mtDNA (Sup. Table 4) was labelled at the 5’;-end with [γ-^32^P]ATP using T4 Polynucleotide kinase (ThermoScientific) according to the manufacturer’s instructions. Probe hybridization was performed using standard procedure. The membrane was exposed to a storage phosphor screen and scanned with the Typhoon 9400 device (Amersham Biosciences).

### POLG DNA extension assay

The exonuclease-deficient catalytic subunit of polymerase γ POLG A and the processivity subunit POLG B were purified as previously described (24). The polymerase γ DNA extension assay was performed using a protocol modified from Jamroskovic et al, 2020 (20). Briefly, 1 μM of TET-labelled annealed template (Sup. Table 4) and pre-incubated POLG A_exo-_ and POLG B (625 nM and 937.5 nM respectively) were mixed in reaction buffer (50 mM Tris HCl (pH 7.6), 10 mM KCl, 50 mM MgCl_2_, 2 mM DTT, 200 ng/μL BSA, 200 μM of dNTPs). When indicated, recBG4 (Millipore Merck) was incubated with the DNA template in reaction buffer at 37 °C for 30 min before starting the reaction. The reactions were carried out at 37 °C for the indicated time points and blocked by the addition of 1:1 stop solution (95 % formamide, 20 mM EDTA (pH 8) and 0.1 % bromophenol blue). Denatured samples were separated over a 10 % polyacrylamide Tris-Borate-EDTA (TBE) gel containing 25 % formamide and 8 M urea. The fluorescent signal was detected with a Typhoon 9400 scanner (Amersham Biosciences). The intensity of each band was quantified as a percentage of total lane signal using Image Quant TL 8.1 software (GE Healthcare Life Sciences). Oligonucleotides used are listed in Sup. Table 4.

### CD spectroscopy

3 μM of each oligo were annealed/folded in 10 mM K-phosphate buffer (pH 7.4) with 100 mM KCl by heating for 5 min at 95 °C and then allowed for cooling to room temperature for overnight. A quartz cuvette with a path length of 10 mm was used for the measurements in a JASCO-720 spectropolarimeter (Jasco Internatiol Co. Ltd.). CD spectra were recorded at 25 °C over λ = 195-400 nm with an interval of 0.2 nm, scan rate of 100 nm/min and 3 times accumulation. Oligonucleotides used are listed in Sup. Table 4.

### EMSA

1 μM oligonucleotides was labelled at the 5’;-end with γ-^32^P-ATP in a reaction catalysed by T4 polynucleotide kinase (PNK) for 60 min at 37 °C. PNK was inactivated by incubating at 65 °C for 10 min. Labelled oligonucleotides were purified by a G50 column (GE Healthcare). 100 nM of 5’;-^32^P end-labelled G4 substrates in 1 mM Tris-HCl (pH 7.5) and 100 mM KCl were incubated at 95 °C for 5 min and then allowed to cool down to room temperature to allow folding into G4 structures.

For the EMSA reaction, 1 nM folded G4 oligonucleotides (or scamble oligonucleotides) was mixed with increasing concentrations of BG4 (0, 1.56, 3.13, 6.25, 12.5, 25, 50 and 100 nM). Reactions (15 μl) containing 1 mM Tris-HCl (pH 7.5), 0.25 mg/mL BSA, 0.1 M KCl, 10 nM MgCl_2_ and 10 % glycerol were incubated at 37 °C for 10 min before separation over a 4.5 % native acrylamide gel at 100 V for 35 min. Gels were dried for 1.5 h at 80 °C before exposure to phosphoimager screen. Bands were visualized using a Typhoon Scanner 9400 and ImageJ Software. Oligonucleotides used are listed in Sup. Table 4. Recombinant MtSSB was purified as previously described (25).

### ChIP

For each treatment, cells were plated into two 15 cm plates two days before harvesting at 1/6 dilution (ca 40 million cells/plate were collected for each experiment). For ρ0 cells, four 15 cm plates were used (ca. 20 million cells/plate). Cells were treated with 100 ng/mL doxycycline for 24 h and, if need, with 0,5 μM of RHSP4 or 200 μM ddC for 3h.

Cross-linking was performed with 1 % Methanol-Formaldehyde in 1x PBS for 10 min directly on the plates, followed by quenching with 125 mM Glycine for 5 min. Cells were collected in ice-cold 1x PBS, washed several times and lysed in 1 mL ChIP lysis buffer (140 mM NaCl, 50mM HEPES-KOH (pH 7.5), 1 mM EDTA, 1% Triton X-100, 0.1% Sodium deoxycholate) supplemented with Halt protease inhibitor cocktail (ThermoFisher Scientific). DNA was sheared with a Covaris E220 instrument to obtain mtDNA fragments of about 300 to 500 bp. The sheared DNA was pre-cleared for 1 h with Pierce Protein A/G Magnetic Beads. 25 μL and 10 μL were collected at this stage as INPUT samples for immunoblot and sequencing analysis respectively. 350 μL of each sample were then incubated overnight with 10 μg of FLAG M2 antibody (Sigma) or mouse IgG (Santa Cruz) followed by 2 h and 30 min incubation with 20 μL Pierce Protein A/G Magnetic Beads. Beads were collected using a magnetic rack and the supernatant was stored for protein analysis (non-bound fraction). Beads were subjected to subsequent washing steps with SDS Buffer (140 mM NaCl, 50 mM HEPES-KOH (pH 7,5), 1 mM EDTA, 0.025 % SDS) two times, High salt wash buffer (1 M NaCl, 50 mM HEPES-KOH (pH 7,5), 1 mM EDTA), TL wash buffer (20 mM Tris HCl (pH 7.5), 250 mM LiCl, 1 mM EDTA, 0.5 % NP-40, 0.5 % Sodium deoxycholate), TE buffer two times. Samples were eluted in TE buffer with 1 % SDS at 65° C for 2 min followed by vortexing. For sequencing and qPCR, samples were reverse cross-linked ON at 65 °C and treated with 100 ng/mL RNaseA for 15 min at 37 °C and 20 μg proteinase K for 2 h at 56 °C to remove RNA and proteins respectively. The samples were finally purified with ChIP DNA clean and concentrator kit (Zymo Research). Samples were quantified by Qbit. ChIP-Seq library preparation and sequencing was performed by Novogen. Briefly, the DNA was subjected to mechanical fragmentation to achieve the proper size for library preparation. The NEBNext Ultra II DNA Library Prep Kit (New Englang Biolab) was used for sequencing library preparation according to the manufacturer’s protocols. The library was quantified by Qubit Fluorometer (ThermoFisher) and real-time PCR, and size distribution was detected with Bioanalyzer (Agilent). Quantified libraries were pooled and sequenced on the Illumina NoveSeq 6000 sequencing platform. The sequencing strategy was pair-end 150bp. Between 40M and 60M reads were generated for each individual sample.

For immunoblot analysis, samples were diluted in 1x Laemmli Sample Buffer (Biorad) with 2.5 % β-mercaptoethanol and denatured before separation on SDS-PAGE as described above. For each sample, 10 μL of immunoprecipitated fraction and 5 μL of INPUT and unbound fraction were separated.

### ChIP-seq data analysis

The quality of the reads was examined using FastQC (0.11.2) (https://www.bioinformatics.babraham.ac.uk/projects/fastqc/). The reads were quality filtered using TrimGalore (0.4.0) (https://www.bioinformatics.babraham.ac.uk/projects/trim_galore/) including reads with a minimum quality of 20 and a minimum length of 30. The Quality filtered reads were mapped towards the human reference genome (GRCh38) using BWA (0.7.5a) (26). To evaluate the quality of the ChIP-seq experiment, i.e. if there is significant clustering of enriched DNA sequence reads at locations bound by BG4, the cross-correlation of each sample was calculated using phantomPeaks (1.1) (27, 28) yielding to NSC values between 1.005558 and 1.042857 and RSC values between 0.8999861 and 1.060696. PCR duplicates were removed using Picard (2.1.0) (https://broadinstitute.github.io/picard/) and artifacts in the genome (blacklist of GRCh38, ENCFF356LFX) were removed using bamUtils (29) from NGSUtils (0.5.9-b4caac3) (30). Enrichment plots were generated with deepTools (2.5.1-1-e071ca1) (31) to assess if the antibody-treatment was sufficiently enriched to separate the ChIP signal from the background signal.

For peak calling, the standard pipeline for ChIP-Seq analysis from the Galaxy platform was followed (32). Briefly, adaptors were removed with Trimmomatic (Galaxy Version 0.36.4), reads were mapped against Human genome hg38 using BWA-MEM (Galaxy Version 0.7.17.1) and non-uniquely mapped reads were removed using SAMtools Filter SAM or BAM (Galaxy Version 1.8+galaxy) selecting a minimum MAPQ quality score of 20. Finally, peak calling for FLAG samples against IgG samples was performed using MACS2 (Galaxy Version 2.1.1.20160309.6). Unless otherwise stated, default parameters were applied in the analysis.

The depth per position and the total amount of high-quality mapped reads (Q20) towards the mitochondria (MT) was calculated using SAMtools (1.9) (33). Only positions with depth higher than 50 were included in further analyses. The samples were normalized for sequencing depth by dividing each position with the total number of mapped reads in the MT for each sample and multiplied by 100. The ratio between the FLAG sample and the INPUT sample was calculated adding a pseudo-count of 1. An inhouse script was used to identify the resulting peaks. Positions with at least 50 contiguous nucleotides with a ratio of at least one were selected. From the summit +/-50 bp were extracted as peak coordinates. Finally, to visualize the peaks coverage, circular plots were generated using Circos (0.69) (34). The linear plots displaying the averages of the sample ratios were generated in R (4.0.1) (https://www.r-project.org/).

### qPCR

2 μL of FLAG or IgG pull down and of a 1:10 dilution of the INPUT were used for amplification using qPCRBIO SyGreen Mix (PCR Biosystem) in a LightCycler96 (Roche) instrument. Oligonucleotides for nuclear and mitochondrial DNA amplification are indicated in Sup. Table 1. Each sample was run in triplicate. Mean Ct values were used to calculate the amount of FLAG and IgG signal with respect to the INPUT. Mean Ct values of the INPUT were corrected for the dilution factor.

## STATISTICAL ANALYSIS

Samples size and number of replicates are indicated for each experiment. Two tail two samples t-test with assumed equal variance was used to determine significant differences. Variance was determined by F test. A p-value < 0,05 was considered significant. All calculations were performed in Microsoft Excel and Origin 2017 software.

## RESULTS

### Establishing a cell line expressing mitochondrial targeted BG4 (mitoBG4)

To detect the formation of G4s specifically in the mtDNA in human cell lines without interfering with the G4s in the nucleus, we took advantage of the BG4 antibody. BG4 is a single chain antibody fragment that can specifically recognize G4s and was previously used to analyze nDNA and RNA G4s in fixed cells (12). We established a cell line expressing a variant of BG4 (mitoBG4) that is targeted to the matrix of mitochondria, where the mtDNA resides. To ensure the correct mitochondrial localization, the BG4 sequence with C-terminal FLAG tag was fused to the mitochondrial targeting signal (MTS) of TFAM (mitochondrial transcription factor A)(18). The obtained MTS-BG4-FLAG sequence was subsequently inserted in the genome of HEK (Human embryonic kidney) 293 cells using the Flp-In T-REx system. This system permits the locus-specific integration of the gene of interest under the control of a doxycycline-inducible promoter, thus allowing the modulation of gene expression (Sup.Fig. 1A).

**Figure 1:**
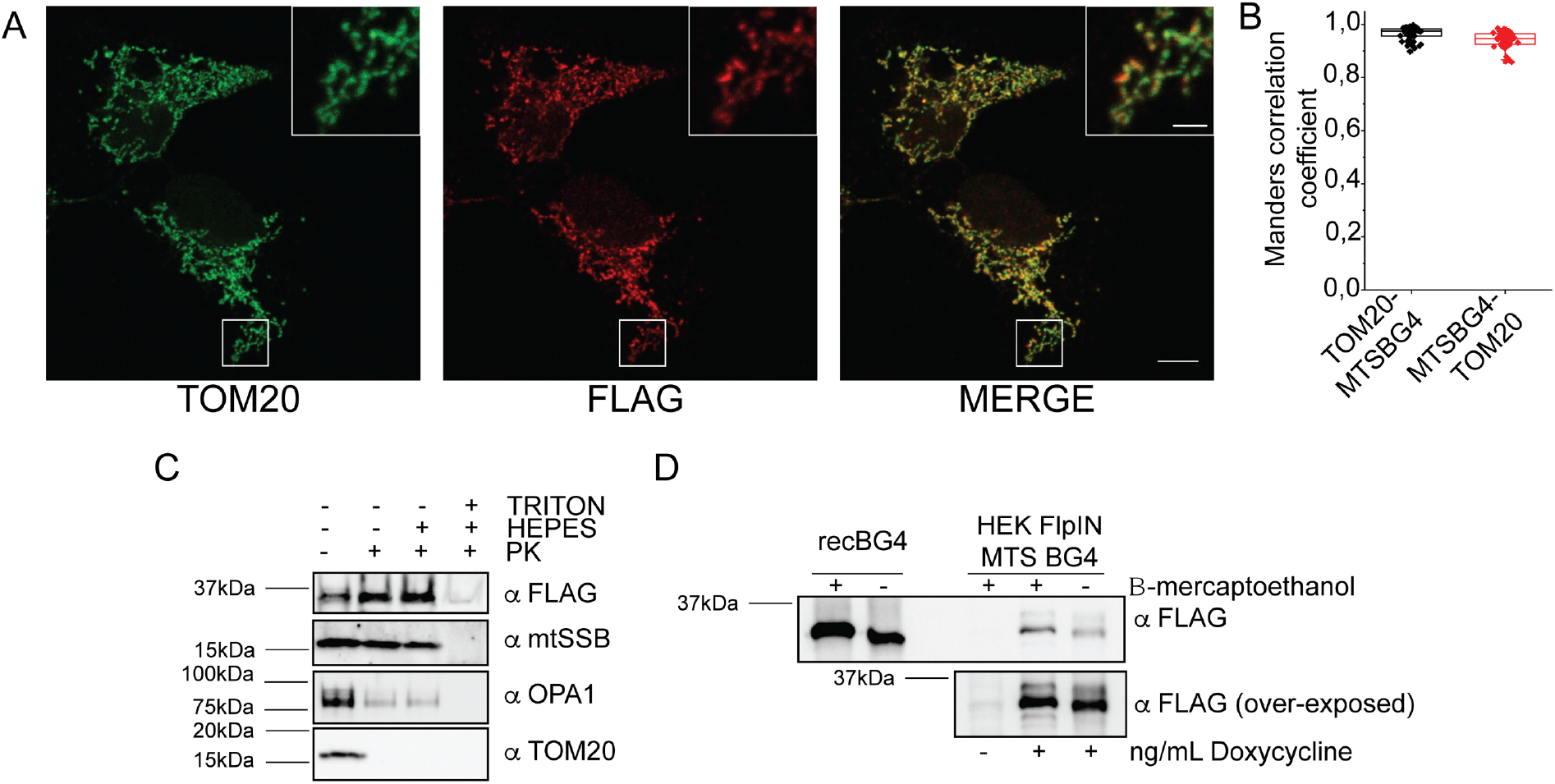
MTS-BG4 is targeted to the mitochondrial matrix and is properly folded. A. Representative immunofluorescence images of induced mitoBG4 cells immunolabeled with FLAG antibody (to detect mitoBG4) and TOM20 (mitochondrial outer membrane marker) antibody. Scale bars: 10 μm and 2 μm in the magnified box. B. Quantification of the reciprocal co-localization of TOM20 and mitoBG4 from experiment in A (50 cells analyzed). C. Mitochondria Proteinase K accessibility assay. Intact mitochondria (Triton – and Hepes -), mitoplasts (Triton – and Hepes +) or detergent solubilized mitochondria (Triton + and Hepes +) from induced mitoBG4 cells were separated on SDS-PAGE. Antibody against mtSSB (mtDNA single strand DNA binding protein), OPA1 (Optic Atrophy 1) and TOM20 were used to detect mitochondrial matrix, inner mitochondrial membrane and outer mitochondrial membrane respectively. D. Immunoblot under native (-B-mercaptoethanol) or reducing conditions (+ B-mercaptoethanol) of recombinant BG4 protein (recBG4) or total cell extract from induced mitoBG4 cells were separated on SDS-PAGE. Samples were probed with FLAG antibody to detect mitoBG4. Non-induced (-doxycyline) mitoBG4 cells were added as control.

Upon induction with doxycycline, the expression of mitoBG4 in hygromycin-B resistant selected cells (i.e. with MTS-BG4-FLAG integrated in the Flp locus) was detected by Western blot using an antibody against the FLAG tag. We detected a time- and dose-dependent expression of mitoBG4 (Sup.Fig. 1B). Two bands were visible, a higher molecular weight band corresponding to mitoBG4 retaining the MTS peptide and a lower band corresponding to mitochondrially imported mitoBG4 with cleaved MTS. The same band pattern is present in TFAM-overexpressing cells (35).

### mitoBG4 localizes to mitochondria and is properly folded

We then looked at the localization and folding of mitoBG4 within the cellular environment. Co-Immunohistochemistry (cIHC) with the mitochondrial marker TOM20 revealed co-localization of mitoBG4 with mitochondria (Fig 1A and 1B), which was further confirmed by a fractionation assay (Sup.Fig. 1C). In addition, a Proteinase K accessibility (PKA) assay demonstrated that mitoBG4 is localized to the mitochondrial matrix, similarly to the mitochondrial single stranded DNA-binding (mtSSB) protein, as it is protected against Proteinase K digestion unless the inner mitochondrial membrane is solubilized (Fig. 1C).

Finally, since BG4 corresponds to the fragment crystallisable region (scFv fragment) of the antibody and requires formation of disulfide bonds to be properly folded, we performed non-reducing SDS-PAGE to test its intracellular conformation. Separation of protein extracts from induced mitoBG4 cells in non-reducing conditions revealed that native mitoBG4, like the recombinant BG4 (recBG4), ran at different size than its reduced counterpart (treated with β-mercaptoethanol), suggesting that mitoBG4 protein forms disulphide bonds when expressed in the cells (Fig. 1D).

Taken together, these data indicate that mitoBG4 localizes and properly folds in the mitochondrial matrix, where the mtDNA also resides.

### mitoBG4 co-localizes with mitochondrial nucleoids

We then set out to analyse whether mitoBG4 interacts with the mitochondrial nucleoids, the nucleoprotein complexes in which the mtDNA is organized. cIHC of induced mitoBG4 cells showed mitoBG4 protein to co-localize with the main mtDNA nucleoid component, TFAM (Fig. 2A and 2C). This co-localization was comparable to that of mitoBG4 and mtSSB, another known nucleoid protein (Fig. 2B and 2C).

**Figure 2:**
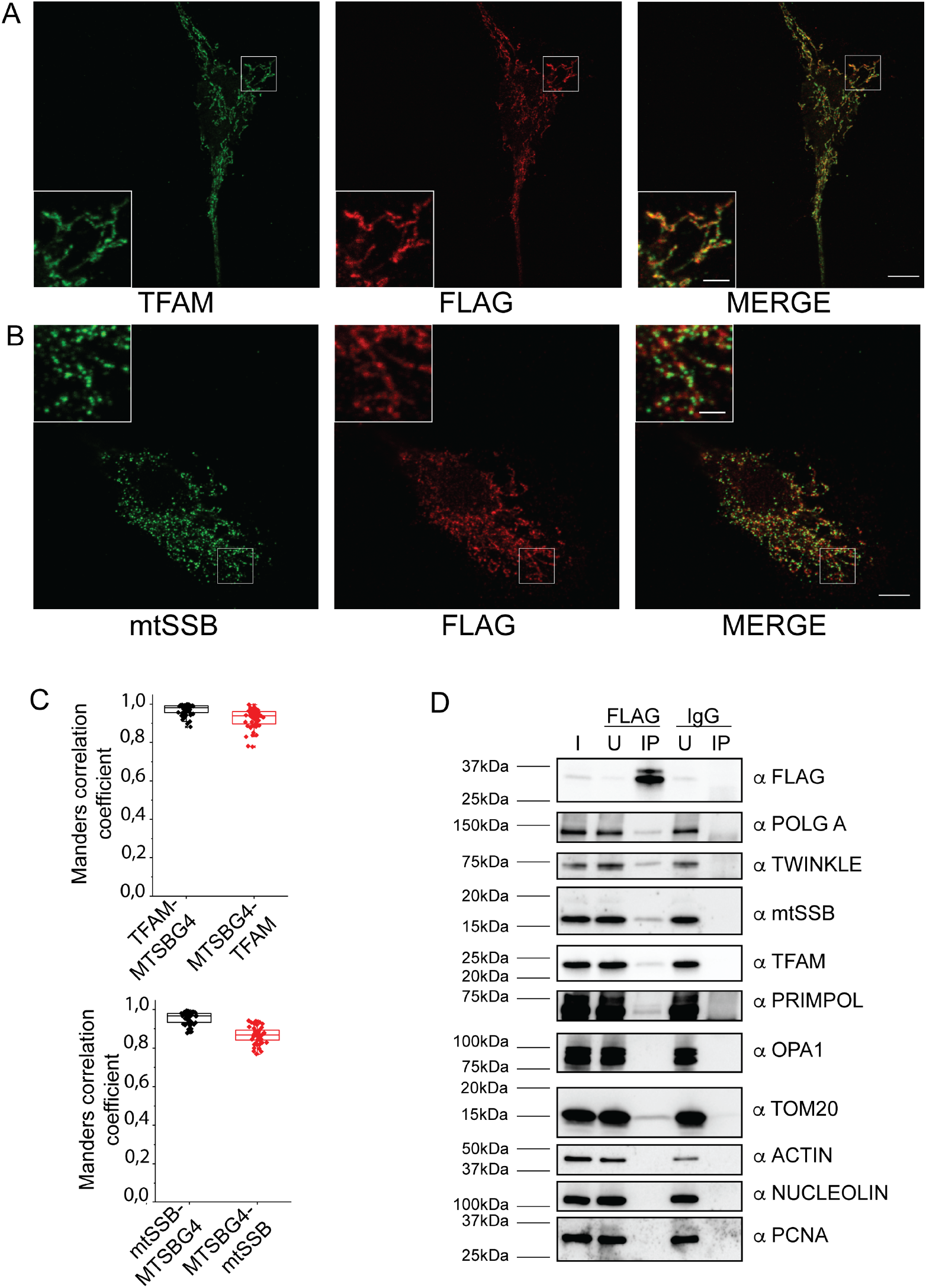
mitoBG4 interacts with mtDNA nucleoids. A and B. Representative immunofluorescence images of induced mitoBG4 cells immunolabeled with FLAG antibody (to detect mitoBG4) and TFAM (A) or mtSSB (B) antibodies. Scale bars: 10 μm and 2 μm in the magnified box. C. Quantification of the reciprocal co-localization of nucleoids marker and mitoBG4 from experiment in A (up-58 cells analyzed) and in B (bottom-48 cells analyzed). D. Immunoblot of Input (I), unbound (U) and pull-down (IP) fractions from FLAG and IgG immunoprecipitation of induced mitoBG4 cells. The blot was probed with the indicated antibodies. POLGA (mitochondrial Polymerase γ, subunit A), mitochondrial helicase TWINKLE are component of the mtDNA replication machinery; PRIMPOL is a primase-polymerase with mitochondrial localization. PCNA and NUCLEOLIN are nDNA binding proteins. ACTIN is a component of the cytoskeleton.

To corroborate these results, we performed FLAG-immunoprecipitation after mitoBG4 expression. While the proteins of the mtDNA replisome and other mtDNA-interacting proteins (PRIMPOL and TFAM) co-precipitated with mitoBG4, we did not detect any interaction with a mitochondrial inner membrane protein (OPA1) nor the cytosolic protein ACTIN or the nuclear proteins PCNA and NUCLEOLIN (a nuclear DNA G4 binding protein (36)) (Fig 2D). Reciprocal immunoprecipitation with antibodies against mtDNA-binding proteins confirmed their interaction with mitoBG4 (Sup.Fig. 2). Notably, TOM20 was also detected in the mitoBG4 IP extract. This protein is required for mitochondrial import and apparently interacts with the non-processed form of mitoBG4 before it reaches the mitochondrial matrix (see Sup.Fig. 2A, note that mainly the higher MW band is detected in the bound fraction upon TOM20 IP).

We conclude that mitoBG4 directly interacts with the mitochondrial nucleoids.

### mitoBG4 does not affect mtDNA maintenance

mitoBG4 expression in cells represents a feasible tool to detect endogenous mtDNA G4s formation only if its induction does not disrupt mtDNA maintenance or modifies G4 structure stability. Recombinant mitochondrial DNA polymerase γ (POLG) has been shown to pause DNA synthesis when encountering G4 structures on the DNA template (11, 37). DNA polymerase stop assays on a previously tested mtDNA G4 template (11) confirmed these results, showing that G4 structures are an obstacle for POLG. However, extended reaction times enabled POLG to partially bypass the G4 structure and synthesize a full-length run-off DNA product (Sup.Fig. 3B lines 4 and 5). The amount of pausing at the G4 structure can therefore be seen as a relative measurement of the G4 structure stability. In agreement with this, chemical stabilization of the G4 structure by PhenDC3 led to a reduction of the run-off DNA product (Sup.Fig. 3B lines 15 to 17). In contrast, addition of recBG4 to the reaction, which binds the G4 structure (Sup.Fig. 3C), did not alter the amount of full-length DNA product (Sup.Fig. 3B compared lines 2 to 5 with 10 to 13). The stalling of POLG is specific for the presence of the G4 structure, as the same reaction performed on a mtDNA sequence that did not contain a putative G4-forming sequence (PGS) did not result in any pausing of POLG (Sup.Fig. 3B, lines 19 to 34). We concluded that BG4 binding to mtDNA is not likely to disrupt DNA synthesis by the major mitochondrial DNA polymerase POLG.

**Figure 3:**
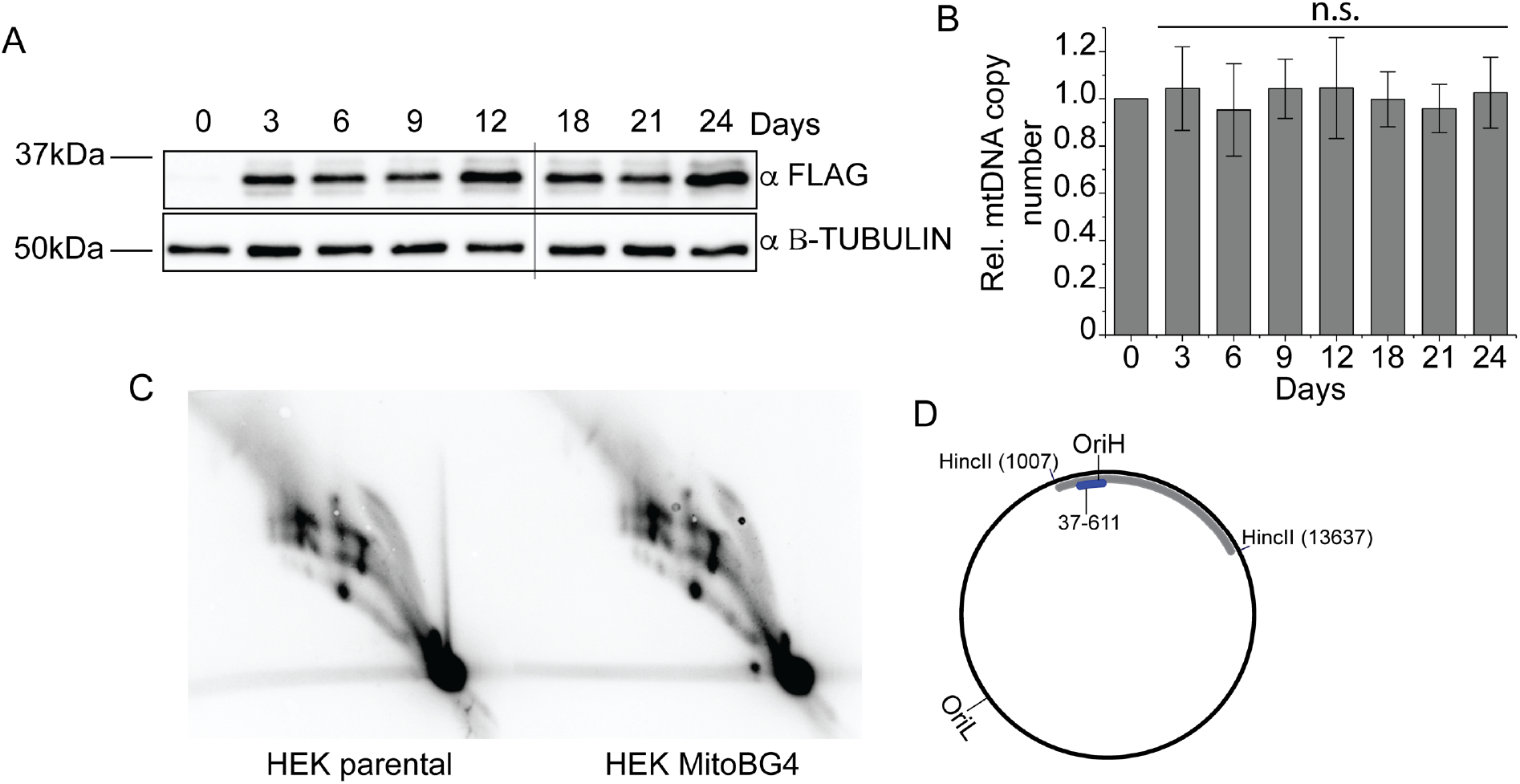
mitoBG4 does not affect mtDNA maintenance. A. Immunoblot of total cells lysate from mitoBG4 cells induced with 10 ng/mL doxycycline for the indicated days. Anti B-TUBULIN was used as loading control. B. mtDNA copy number of mitoBG4 cells induced with 10 ng/mL doxycycline for the indicated days. MtDNA copy number was determined by qPCR from total DNA. The relative mtDNA copy number was expressed as the ratio between mtDNA D-loop region and nuclear DNA B2M region and normalized to untreated samples. Data represent mean ± s.d. of three independent experiments. Analysis of the data was performed with two-tail two samples t-test. n.s. = not significant. C. 2D-AGE of HincII-digested mtDNA from parental HEK Flp-In T-REx cells transfected with the empty vector pcDNA5 (HEK parental) and induced HEK mitoBG4 cells (HEK MitoBG4), probed for the OriH-containing fragment as illustrated in D. No differences in the replication intermediates can be observed. D. A schematic illustration of human mtDNA showing the HincII restriction sites used for linearization of mtDNA. The probe used for Southern blot hybridization is indicated in blue.

We then explored the effect of mitoBG4 expression on mtDNA in the mitoBG4 cells. Long-term treatment with doxycycline (up to 24 days) did not have any effect on the mtDNA copy number (Fig. 3A and B). MtDNA copy number analysis might, however, not be able to detect mild interference of the mtDNA replication process. To address this more specifically, we used two-dimensional agarose gel electrophoresis (2D-AGE) analysis of the mitochondrial DNA replication intermediates (19). The doxycycline induction of mitoBG4 did not alter the pattern of mtDNA replication intermediates compared to control cells (Fig. 3C and D).

We conclude that BG4 does not interfere with *in vitro* G4-stability and that its intracellular expression has no effect on the maintenance of mtDNA. This system can thus be used to detect the regions of mtDNA in which G4 structures are formed in human cells without affecting mtDNA metabolism.

### mtG4-ChIP enriches for mtDNA G4 containing regions

We then set up a ChIP protocol to map the formation of G4s in the mitoBG4 cells by inducing the expression of mitoBG4 protein and subsequently perform a pull down with the FLAG antibody (mtG4-ChIP). To avoid alteration of the mitoBG4 binding to G4-DNA during the time of mitochondrial extraction (which can take up to several hours), we performed a cross-linking step on intact cells and sheared total genomic DNA (nDNA and mtDNA) from total cell extract (Sup.Fig. 4A). We ensured the proper shearing of the mtDNA (fragments size between 500-300bp) using Southern blot against a mtDNA specific probe (ND5) (Sup.Fig. 4B). The FLAG pulldown extract and the INPUT samples were subsequently sent for next generation sequencing (NGS) using Illumina Platform.

**Figure 4:**
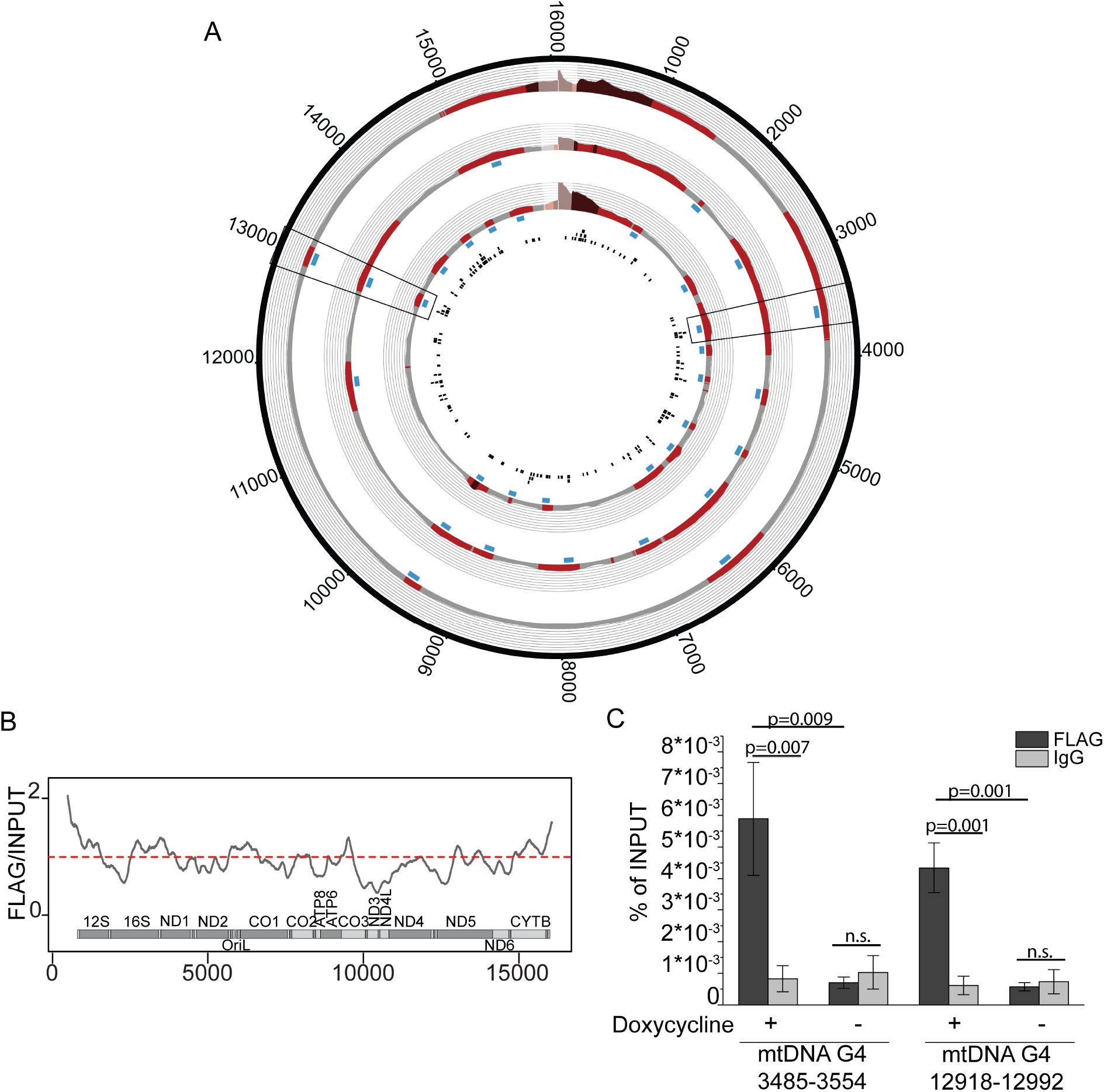
mtG4-ChIP enriches for mtDNA G4s. A. mtG4-ChIP-seq profile for induced mitoBG4 cells pulled down with FLAG antibody. FLAG signal was normalized to the INPUT sample and expressed as ratio of FLAG vs INPUT. Areas in red indicate enrichment over INPUT samples (gray <1, light red 1 > x > 1,5 and dark red > 1,5). The blue boxes beneath the plots are the narrow peaks. The black boxes in the inner circle are predicted G4 sequences (using the G4Hunter algorithm). Three biological replicates are shown. D-loop region (in lighter color) was excluded from the analysis. The black boxes show the regions analyzed by ChIP-qPCR B. Linear plots showing the average FLAG to INPUT signal from the three replicates in figure 4A. MtDNA sequence from nt 500 to nt 16 000 is displayed. C. ChIP-qPCR upon pull down with FLAG antibody and IgG in induced (+) or non-induced (-) mitoBG4 cells. Data are expressed as % of INPUT pull down. Data represent mean ± s.d. of three independent experiments. Analysis of the data was performed using with two-tail two samples t-test. n.s. = not significant.

We initially used a standard method for ChIP-seq analysis (32). However, we did not detect individual peaks for most of the samples (data not shown) likely because the Peak Calling MACS2 software was developed for nDNA analysis and might not take into consideration the multicopy nature and the relatively small size of the mitochondrial genome. Therefore, in order to improve the visualization, we used a manual peak-calling methodology. Sequence analysis upon normalization showed several mtDNA regions enriched with respect to the INPUT (Fig. 4A). The D-loop region was also enriched but was excluded from our analysis as this area was shown to be prone to artefacts in ChIP-seq protocols, as testified by enrichment upon pull down with IgG (38) or with uniquely localized nuclear transcription factors (39). We next performed mtG4-ChIP followed by qPCR (mtG4-ChIP-qPCR) and amplified two mtDNA regions that are enriched in all three replicates (Fig. 4A and 4B). We detected 7-fold enrichment of these two regions compared to IgG, while no enrichment was detected for the cells that were not induced with doxycycline (i.e. no mitoBG4 expression) (Fig. 4C). Analysis of a nDNA region showed no enrichment (Sup.Fig. 5A). We also analyzed by mtG4-ChIP-qPCR two mtDNA regions that were not enriched in the ChIP-seq protocol, and we detected increased signal in the FLAG samples from induced cells, albeit to a lesser extent compared to the ChIP-seq enriched mtDNA regions (Sup.Fig. 5A). However, as the PGS are spread out all over the mtDNA sequence (Fig. 4A) and fragmentation of the mtDNA for the ChIP is always incomplete, also the less abundant G4 structures will increase the overall mtDNA co-purification from mitoG4 expressing cells.

**Figure 5:**
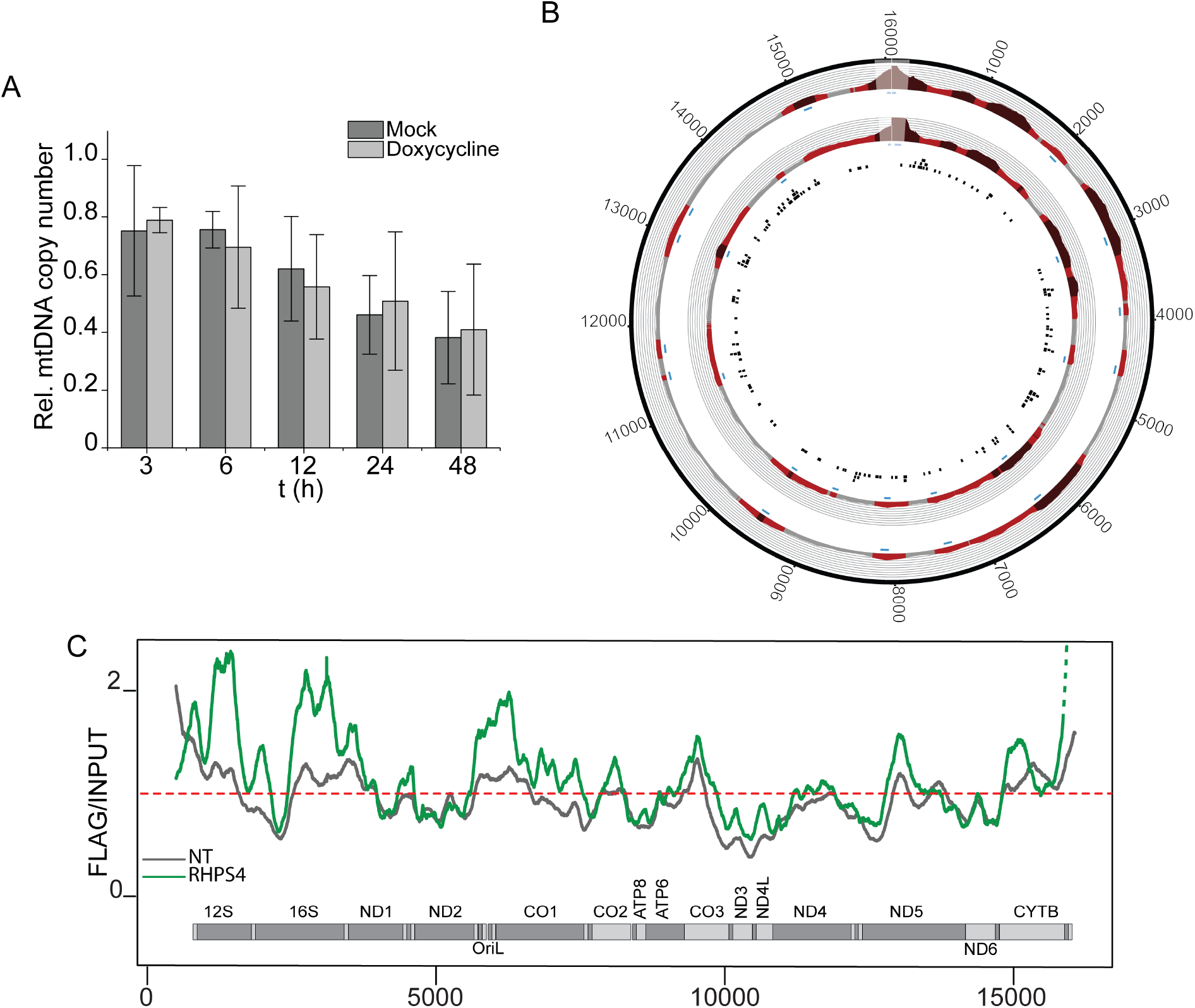
Stabilization of mtG4 causes increased mtG4-ChIP enrichment. A. mtDNA copy number of induced mitoBG4 cells treated with 0.5 μM RHPS4 for the indicated time points. MtDNA copy number was determined by qPCR from total DNA. The relative mtDNA copy number was expressed as the ratio between mtDNA D-loop region and nuclear DNA B2M region and normalized to untreated samples. Data represent mean ± s.d. of three independent experiments. B. mtG4-ChIP-seq profile for induced mitoBG4 cells treated with 0.5 μM RHPS4 for 3 h before pulling down with FLAG antibody. FLAG signal was normalized to the INPUT sample and expressed as ratio of FLAG vs INPUT. Areas in red indicate enrichment over INPUT samples (gray <1, light red 1 > x > 1,5 and dark red > 1,5). The blue boxes beneath the plots indicates the narrow peaks. The black boxes in the inner circle are predicted G4 sequences (using the G4Hunter algorithm). Two technical replicates are shown. C. Linear plots showing the average FLAG to INPUT signal from the doxycycline induced replicates (in grey) and ddC treated replicates (in red). MtDNA sequence from nt 500 to nt 16 000 is displayed.

We selected four PGS, predicted with G4HUNTER, located in the enriched regions in all three replicates and mapping within or in close proximity (< 10nt) from the peak summit detected by MACS2 (Sup.Table 1). Circular dichroism (CD) studies revealed that the selected PGS can fold into parallel G4 structures (positive ellipticity peak at 264nm). By contrast, scramble sequences with non-contiguous G-tracts did not display CD spectra indicative of any G4 topology (Sup.Fig. 6 and Sup. Table 3). We then used electrophoresis gel mobility assay (EMSA) to evaluate the ability of recBG4 to bind these G4 structures *in vitro*. As expected, recBG4 bound to the G4 structures but was unable to bind to a mtDNA sequence that does not contain PGS (Sup.Fig. 6). Taken together, these data suggest that mtDNA G4s are present in human cells under normal growth conditions and that mitoG4-ChIP can be applied for their detection. However, the rather low level of G4 structures detected in mtDNA suggests that mtDNA G4s are transiently formed under normal cellular growth conditions.

**Figure 6:**
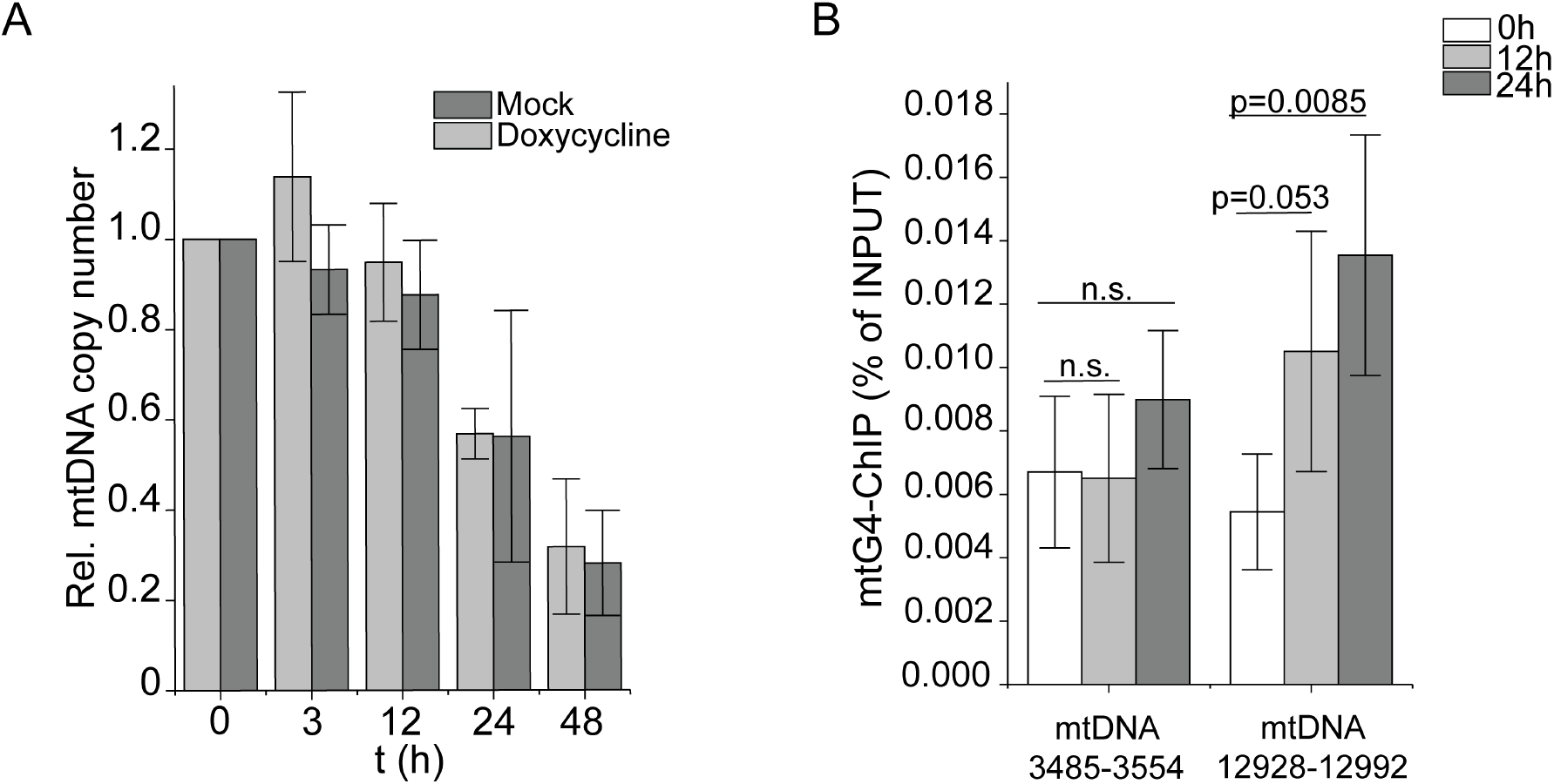
Effect of stalling of the mtDNA machinery in mtDNA G4 formation. A. mtDNA copy number of induced mitoBG4 cells treated with 200 μM ddC for the indicated time points. MtDNA copy number was determined by qPCR from total DNA. The relative mtDNA copy number was expressed as the ratio between mtDNA D-loop region and nuclear DNA B2M region and normalized to untreated samples. Data represent mean ± s.d. of three independent experiments. B. ChIP-qPCR upon pull down with FLAG antibodies in induced mitoBG4 cells treated with 200 μM ddC for the indicated time points. Data are expressed as % of INPUT pull down. Data represent mean ± s.d. of four independent experiments. Analysis of the data was performed using the with two-tail two samples t-test. n.s. = not significant.

### mtG4-ChIP is specific for mtDNA

The nDNA is rich in sequences with high degree of homology with the mtDNA, the so called NUMTs (nuclear DNA mitochondrial sequences) (40). To confirm that the signal detected by mitoG4-ChIPseq protocol is specific for mtDNA and not the result of NUMTs pulldown, we depleted the mtDNA in the mitoBG4 cells by long-term ethidium bromide treatment (41). These cells are essentially ρ0, lacking all mtDNA, as measured by qPCR amplifying two distinct regions of the mtDNA (Sup.Fig. 5B). We then performed the mitoG4-ChIP-seq protocol on the mitoBG4 ρ0 cells lines and detected no residual signal in the mtDNA upon pull down with the FLAG antibody (Sup.Fig. 5C), despite the fact that mitoBG4 was correctly processed and immunoprecipitated (Sup.Fig. 5D). These results confirm that the signal detected in the normal cells by mitoG4-ChIP-seq is specific for the mtDNA.

### Stabilization of mtDNA G4 interferes with mtDNA replication

Next, we investigated the consequence of G4 structures on the progression of the mtDNA replication fork. For this purpose, we took advantage of the known G4 stabilizer RHPS4 described to localize with mtDNA and hypothesized to stabilize G4s in the mtDNA (42). We treated cells with 0.5 μM RHPS4 and performed a time-dependent measurement of the mtDNA copy number. A clear decrease of mtDNA was observed in the presence of the G4-stabilizing compound compared to untreated cells, and after 24 hours mtDNA levels almost halved (Fig. 5A), showing that RHPS4 essentially blocks the progression of mtDNA replication. In accordance, 2D-AGE analysis showed mtDNA replication intermediates accumulation indicative of enhanced replication stalling (Sup.Fig. 7).

**Figure 7.**
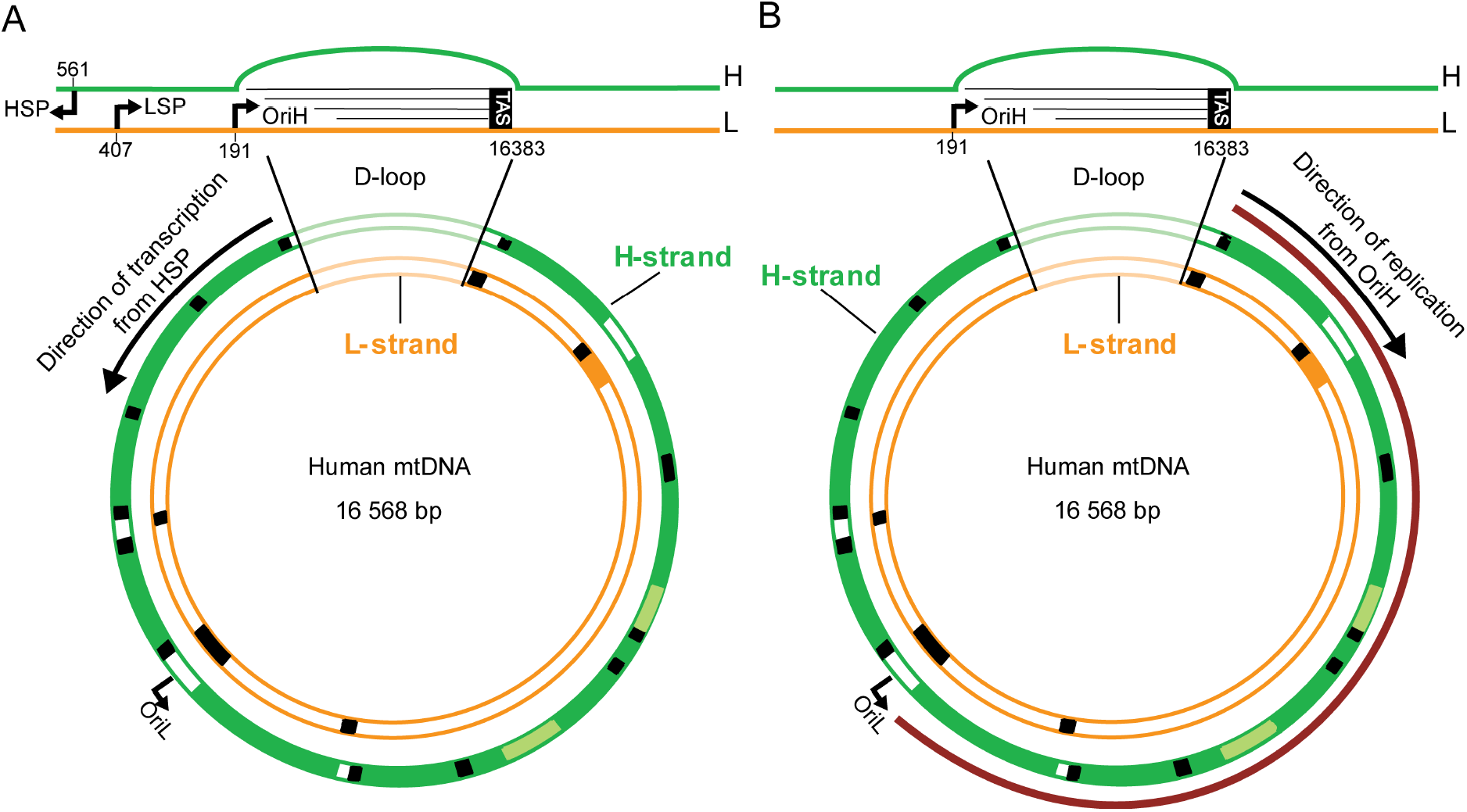
Schematic representation of the human mtDNA including the direction of mtDNA transcription (A) and replication (B). mtDNA is represented anti-clockwise. mtDNA transcription of the H-strand occurs in anti-clockwise direction (black arrow in A). mtDNA DNA replication of the H-strand occurs in clockwise direction (black arrow in B). The control region (D-loop) is depicted in more details. Green: heavy strand (H-strand); orange: light strand (L-strand); red: major arc. OriH and OriL are the origins of replication for heavy and light strand respectively. HSP: H-strand promoter for transcription; LSP: L-strand promoter for transcription. TAS: termination associated-sequences.

To show that the loss of mtDNA is a direct cause of enhanced G4 formation, we performed the mtG4-ChIPseq protocol upon RHPS4 treatment. We treated induced mitoBG4 cells with 0.5 μM RHPS4 for 3 hours, a time where mtDNA copy number was not yet substantially affected (Fig. 5A). Analysis of the sequencing results revealed an overall strong increase in mtDNA sequence reads containing putative G4 sequences upon RHPS4 treatment (Fig. 5B and 5C and Sup.Table 3), indicating that RHPS4 causes stabilization of mtDNA G4s. We conclude that enhanced G4 formation blocks the progression of the mtDNA replication machinery and that the mtG4-ChIPseq is a suitable method to detect an increase in mtDNA G4 formation in human cell lines.

### Stalling of mtDNA replication enhances G-quadruplex formation in cultured human cells

We next asked whether G4 formation was altered when we disrupt the mtDNA replication process. The partially single-stranded nature of mtDNA replication intermediates could make them potentially vulnerable to secondary DNA structure formation (*e.g.* G4s). This might be particularly the case when replication moves slowly, leaving the GC-rich heavy strand exposed for longer time periods (43). To test whether impaired mtDNA replication has any impact on mtDNA G4s-formation, we used the chain-terminating nucleoside analogue 2ʹ-3ʹ-dideoxycytidine (ddC) that is specifically incorporated into mtDNA by POLG and results in mtDNA replication blockage due to chain termination (44). We induced the expression of mitoBG4 in combination with the addition of ddC using conditions (ddC, 200 μM and 3 h) that ensures minimal mtDNA copy number loss (Fig. 6A) but also apparent accumulation of stalled mtDNA replication forks (19), and performed the mtG4-ChIP-seq (Sup.Fig. 8A). Compared to normal growth condition, we detected an increase in the enrichment of some of the mtDNA G4 regions in ddC-treated cells (Sup.Fig.8B and Sup.Table 3). This modest ddC-induced increase of the mtG4-ChIP-seq signal suggests that slowed mtDNA replication might account for enhanced G4 formation on the mitochondrial genome.

To increase the amount of mtDNA molecules that show replication stalling, we tested the effect of longer ddC treatment and performed mtG4-ChIP-qPCR in mitoBG4-expressing cells (12 h and 24 h). Interestingly the ddC-dependent increase of G4 detection after prolonged treatment was more pronounced on the major arc region (12 928-12 992 nt), compared to the minor arc region (3845-3554 nt) (Fig. 6B). These data are consistent with enhanced G4 formation on the exposed parental DNA strand due to a slow progressing mtDNA replication machinery (stalling) during strand-asynchronous mtDNA replication.

## DISCUSSION

In this study we developed a cell model that allows the detection and mapping of G4s specifically in the mtDNA of human cell lines, by targeting the G4-binding synthetic antibody BG4 to mitochondria. We proved that mitoBG4 is correctly localized and folded within the mitochondrial matrix. Moreover, inside the organelle, mitoBG4 interacts with the mtDNA, but does not interfere with the process of mtDNA replication. We subsequently developed a chromatin immunoprecipitation approach specific for mtDNA (mtG4-ChIP) coupled with sequencing that allowed us to detect G4 formation in human cells. The mtG4-ChIP-seq enriches mtDNA sequences that have the ability to form G4s *in vitro*, but also demonstrates that the level of mtDNA-G4 structures is low under regular cell culture conditions, which is in line with the idea that G4s on the mtDNA are transient in nature. To further validate our approach, we employed RHPS4, a well-recognized G4-stabilizing compound, and detected substantial increase of mitochondrial G4 formation in cells after its addition. Furthermore, this increase in mitochondrial G4s leads to a rapid loss of mtDNA, indicating that elevate levels of G4s block the progression of the mtDNA replication fork. In addition, ddC-induced mtDNA replication stalling increases G4 detection by mtG4-ChIP, particularly in the major arc of the mtDNA, suggesting that a slower mtDNA duplication process enhances G4 formation specifically in the region of the mitochondrial genome where mtDNA deletions accumulate in disease.

Human mtDNA, despite being small in size, is highly enriched in putative G4-forming sequences, with a frequency of 6.6 G4s for every 1000 nts, as detected using the G4Hunter software (45). However, the amount of mtDNA G4s detected under normal cell culture conditions is limited. This could be explained by the fact that the formation of G4s in the mtDNA under physiological conditions is prevented by the function of the mtDNA replisome (and the coating by mtSSB) and/or by the action of specific proteins (e.g. the helicases PIF1 and REQL4 or the nuclease helicase DNA2 (46)) in charge of their clearance.

These experimental findings, implying limited formation of mtDNA G4 structures and the absence of specific hotspots for their formation, reflect the observations in patients with mtDNA deletions accumulation. Indeed, most patients, while carrying the genetic defects from birth, usually present with the first symptoms of the disease in adulthood, indicating that the accumulation of mtDNA deletions is a slow process rather than an acute event. In addition, the pattern of deletions found in these patients is extremely variable, and it does not only differ between different patients, but also in different tissues within the same patient (47, 48).

Under regular cell growth conditions, the formation of mtDNA G4s, as detected by mtG4-ChIP-seq, is more prevalent in the region between 0 and 6000 nts, and this is further increased upon treatment with the G4 stabilizer (RHSP4). These data correlate with previous transcriptomic analysis on RHSP4-treated cells presenting with a progressive decrease of transcripts originating from HSP promoters (42). Since these transcripts are transcribed from the H-strand, this observation is consistent with the increased amount of G4 formation that we detect in the H-strand downstream of HSP (Fig. 7A). Thus, under normal cellular conditions, G4s might transiently form in the mtDNA during mitochondrial transcription, but their short-lived presence may not affect the ability of the mitochondrial replication fork to duplicate the mtDNA. However, the persistence of these structures by the interaction with a G4 stabilizer (RHSP4) will lead to reduced transcription and subsequent block of mtDNA replication, resulting in genetic instability. In agreement with this, Butler and co-workers demonstrated that DNA synthesis by two known mitochondrial DNA polymerases (POLG and PRIMPOL) *in vitro* was strongly blocked by stable G4 structures (37).

By contrast, stalling of the DNA replisome induced by the POLG-specific inhibitor ddC revealed enhanced formation of G4 in the major arc of the mtDNA. Human mtDNA replication occurs in a highly strand-asynchronous manner such that the lagging strand (L-strand) is synthesized with considerable delay. The implication of this mode of replication is that the parental H-strand is exposed in single-stranded form for an extended period of time (Fig.7B). The single-stranded nature of the parental H-strand could make it more prone to G4 formation, in particular when replication moves slowly, so that the major arc of the parental H-strand is exposed even longer than under normal conditions. The slower movement of the replication machinery could be due to the presence of ddC, as in Fig. 6B, or to defects in replisome components, as we and others have previously shown that a defective mitochondrial replication machinery can cause slowing of the replication process (19,49,50). This could also explain why most mtDNA deletion breakpoints in patients have been mapped within the major arc, where the replication machinery will more likely be challenged by G4 formation. The G4s formed on the exposed parental H-strand will not hamper nascent H-strand synthesis that uses the parental L-strand as template, but the structures will be encountered during the synthesis of the nascent L-strand. This is compatible with the recently proposed mechanism of mtDNA deletion formation through copy-choice recombination during L-strand synthesis (51). In short, G4s can form a replication barrier, causing dissociation of POLG during synthesis of the nascent L-strand. DNA breathing could allow re-annealing of the 3’ end of the nascent L-strand DNA at a complementary downstream sequence on the parental H-strand, explaining why G4-forming sequences are enriched at mtDNA deletion breakpoints.

Overall, our data and work from others indicate that while G4-forming sequences evolved in the mtDNA genome with a role in regulating transcription and translation (5–7), they do not represent a threat for the progression of the mtDNA replication machinery under normal conditions. However, they could potentially have a negative impact on mtDNA stability when other factors interfere with the process of mtDNA replication.

Taken together, these data indicate that we can now visualize mtDNA G4s formed in cultured human cells without perturbing their dynamics. The mtG4-ChIP protocol will be instrumental in future research to identify the factors affecting G4 formation in mtDNA, the helicases and nucleases involved in their resolution and, ultimately, to understand the mechanistic role of G4s in the generation of pathogenic mtDNA deletions.

## AVAILABILITY

The ChIP-seq data have been deposited to the Sequence Read Archive (SRA) at NCBI (https://www.ncbi.nlm.nih.gov/sra) under the accession number PRJNA811445.

## SUPPLEMENTARY DATA

Supplementary Data are provided in a separate .pdf file.

## ACKNOWLEDGEMENTS

We acknowledge Irene Martinez Carrasco and the Biochemical Imaging Centre at Umeå University and the National Microscopy infrastructure, NMI [VR-RFI 2019-00217], for providing assistance. We thank Shankar Balasubramanian for providing the pSANG10-3F-BG4 plasmid. We thank Paulina Wanrooij, Josefin Forslund and Gorazd Stojkovic for the helpful discussion. We finally acknowledge Paolo Lorenzon for helping in figures preparation.

---

**Sup.Figure 1:**
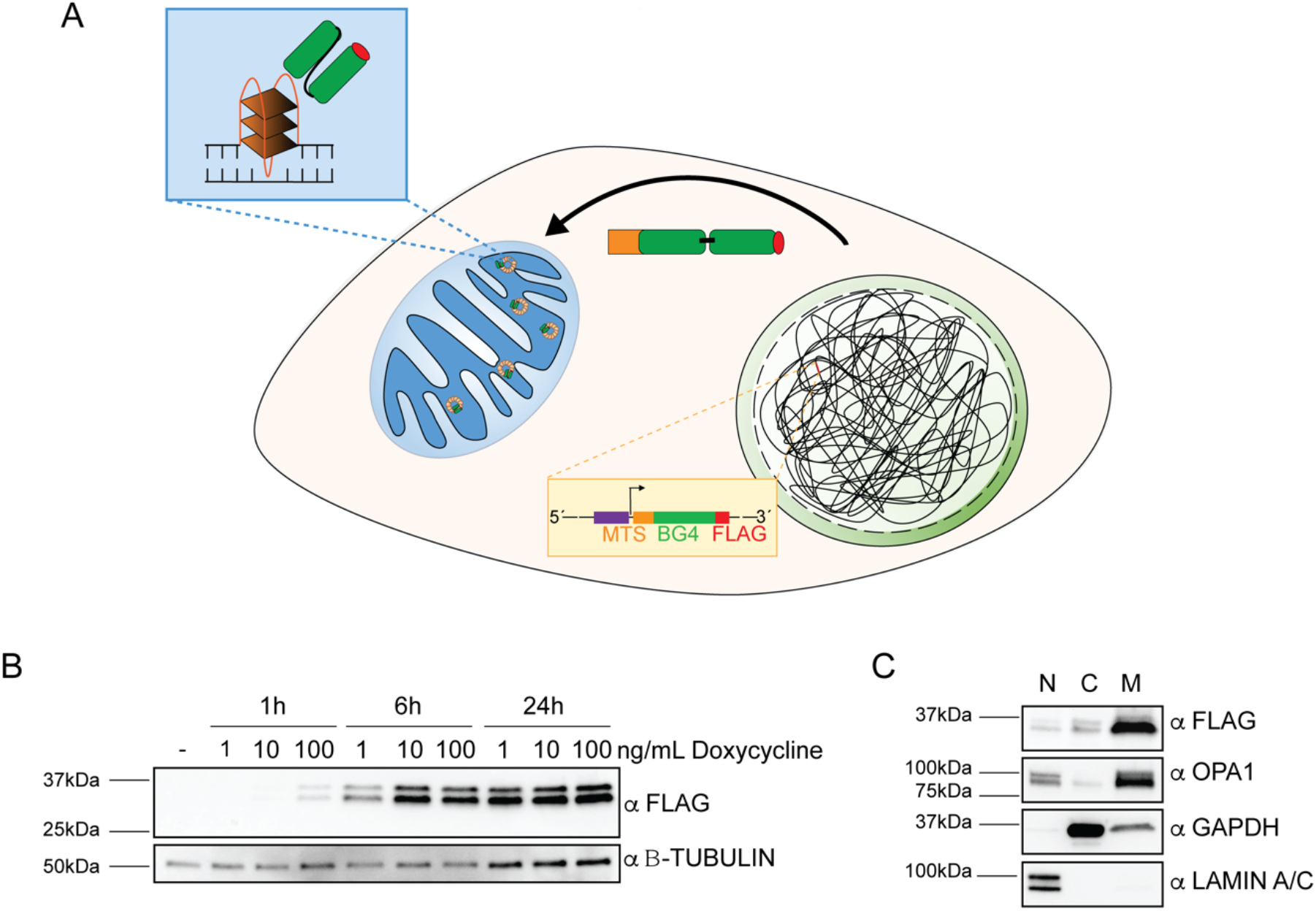
A. Schematic representation of the cell model employed in the study. Briefly, MTS-BG4-FLAG gene is integrated in the genome of the HEK Flp-In T-REx system under the control of an inducible promoter. Upon doxycycline treatment the protein is expressed and targeted to the mitochondrial matrix where, upon cleavage of the mitochondrial targeting sequence and proper folding, can bind to G4 structures in the mtDNA. B. Immunoblot of total cell extracts from mitoBG4 cells treated with different concentrations of doxycycline for the indicated period of time. Samples were probed with FLAG antibody to detect mitoBG4. B-tubulin antibody was used as loading control. C. Immunoblot of nuclear (N), cytosolic (C) and mitochondrial (M) fractions from induced mitoBG4 cells. Antibodies against LAMIN A/C (nucleus), GAPDH (cytosol) and OPA1 (mitochondria) were used to detect the purity of the different fractions.

**Sup.Figure 2:**
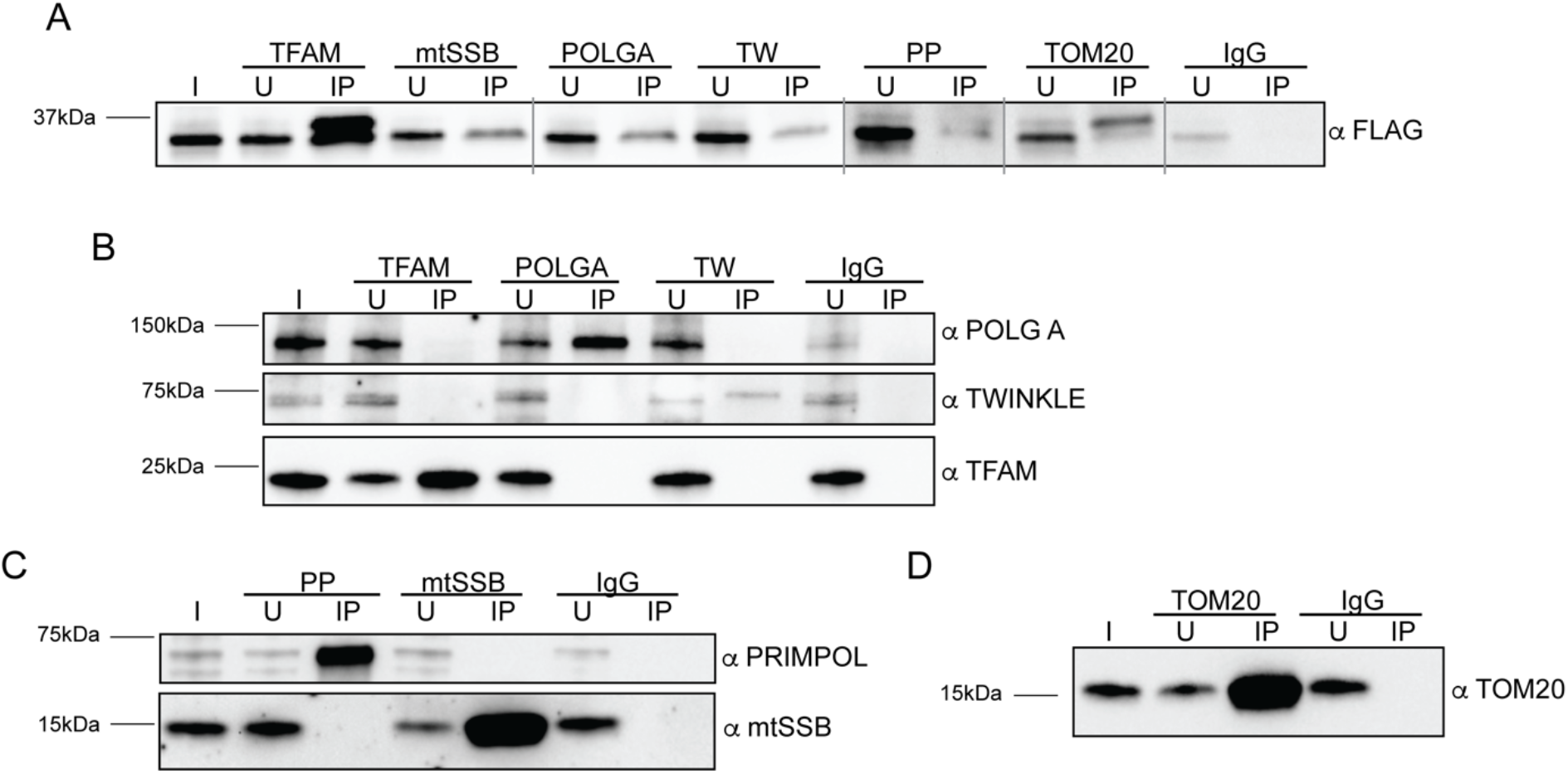
A. Immunoblot of INPUT (I), unbound (U) and pull-down (IP) fractions from immunoprecipitation with the indicated antibodies of induced mitoBG4 cells. The blot was probed with the FLAG antibody to detect mitoBG4. B, C and D. Immunoblot of INPUT (I), unbound (U) and pull-down (IP) fractions from immunoprecipitation with the indicated antibodies of induced mitoBG4 cells.

**Sup.Figure 3:**
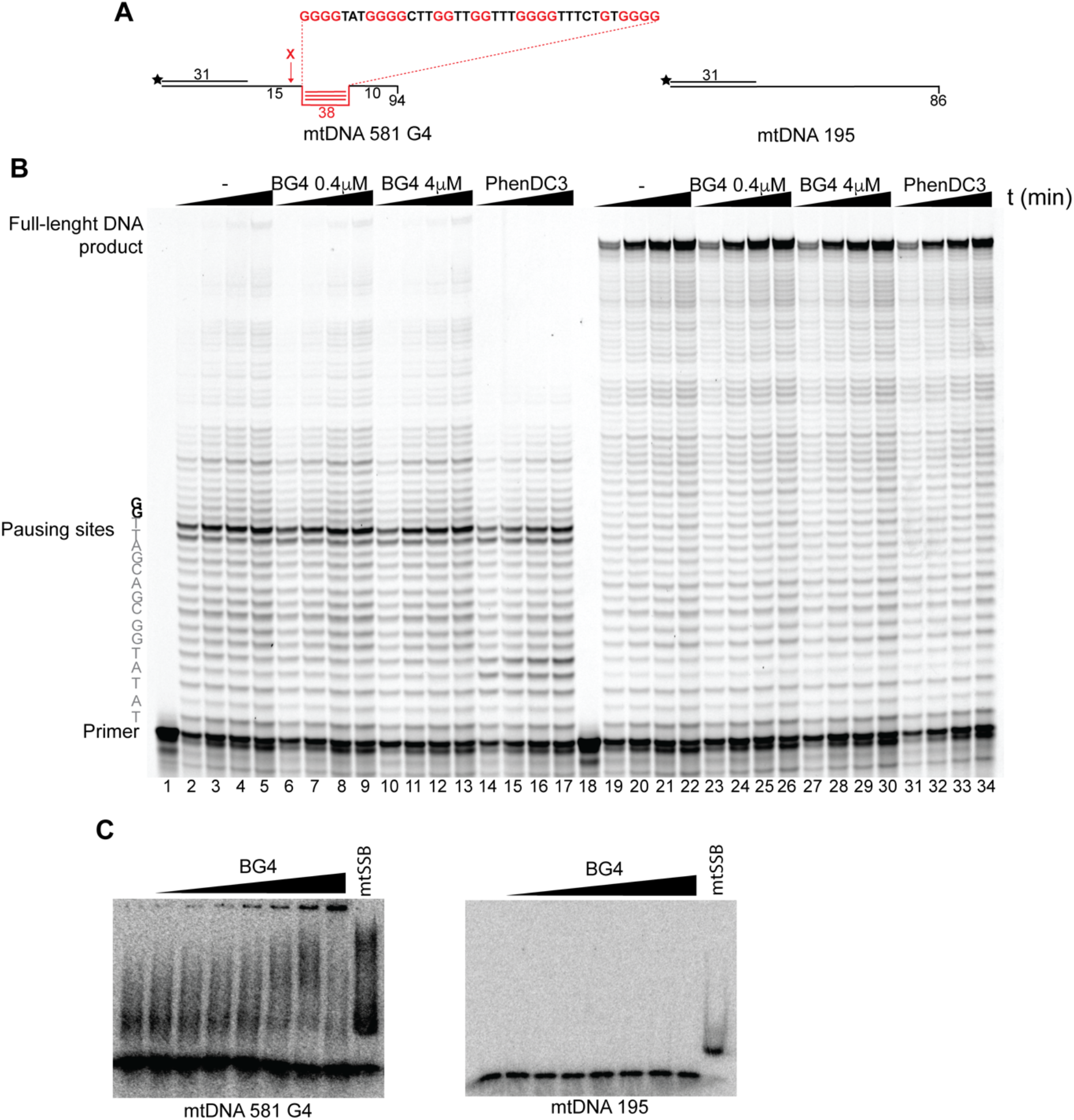
A. Schematic representation of the templates used for the polymerase stop assay. B. Polymerase stop assay on a template with a G4 structures in the presence of increasing concentration of recombinant BG4 (BG4). For each concentration, the reaction was blocked at increasing time points. The template sequence before and around the pausing site are highlighted on the right side. Guanines involved in the G4 formation are indicated in bold. C. Electrophoretic mobility shift assay (EMSA) in the presence of increasing concentrations of BG4 of the oligos used for the Polymerase stop assay. DNA concentration was 1 nM. Recombinant mtSSB (mtSSB) was added as binding control.

**Sup.Figure 4:**
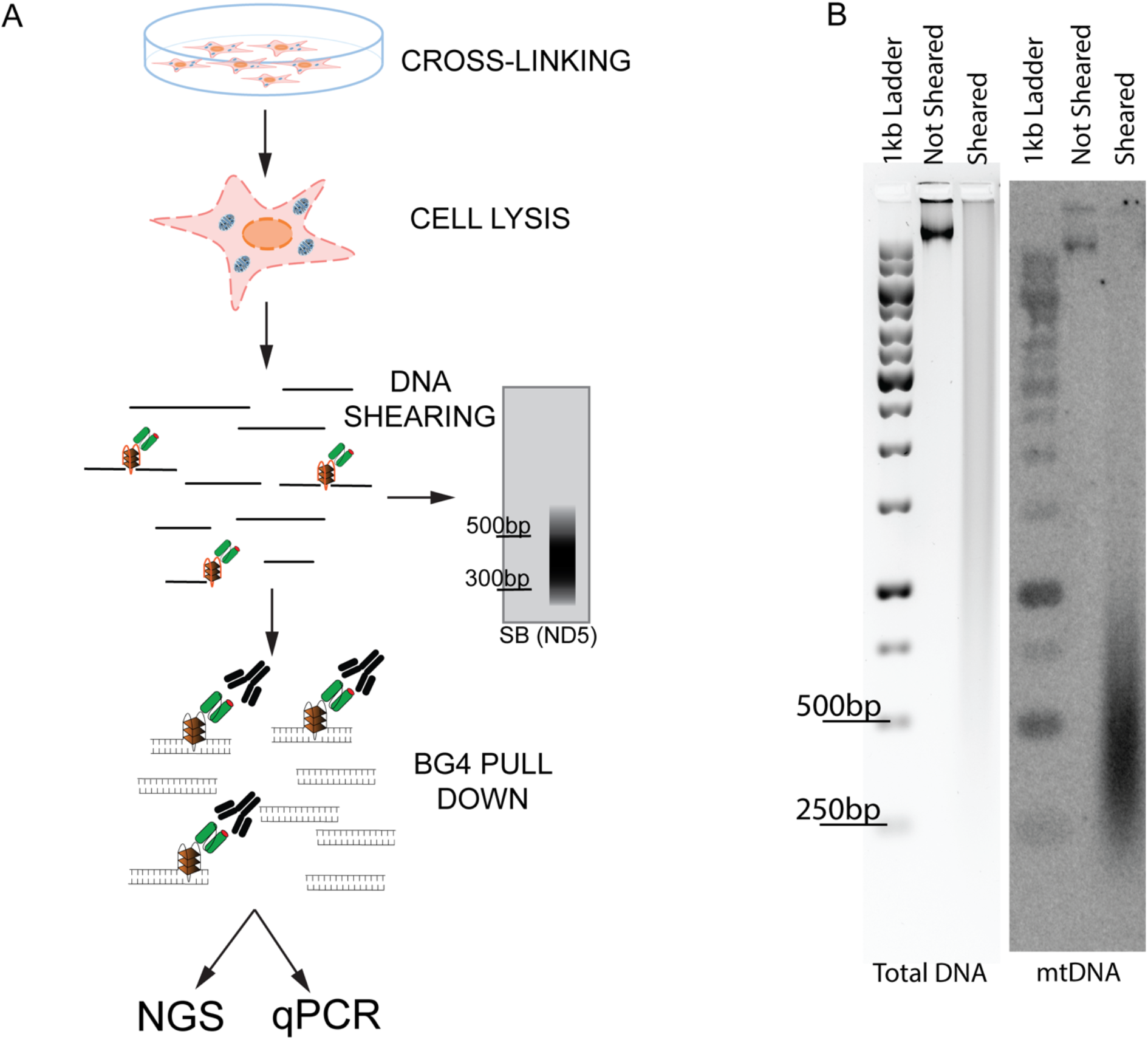
A. Schematic representation of the mtG4-ChIP protocol. B. Southern blot analysis of DNA shearing for the mtG4-ChIP. Reverse-cross linked DNA intact and sheared was separated on agarose gel and detected under UV light upon Ethidium bromide staining to detect total DNA signal (left picture-total DNA). The separated DNA was then blotted on nylon membrane and the mtDNA was detected with a ^32^P-labelled mtDNA specific probe (against ND5 region, right picture-mtDNA).

**Sup.Figure 5:**
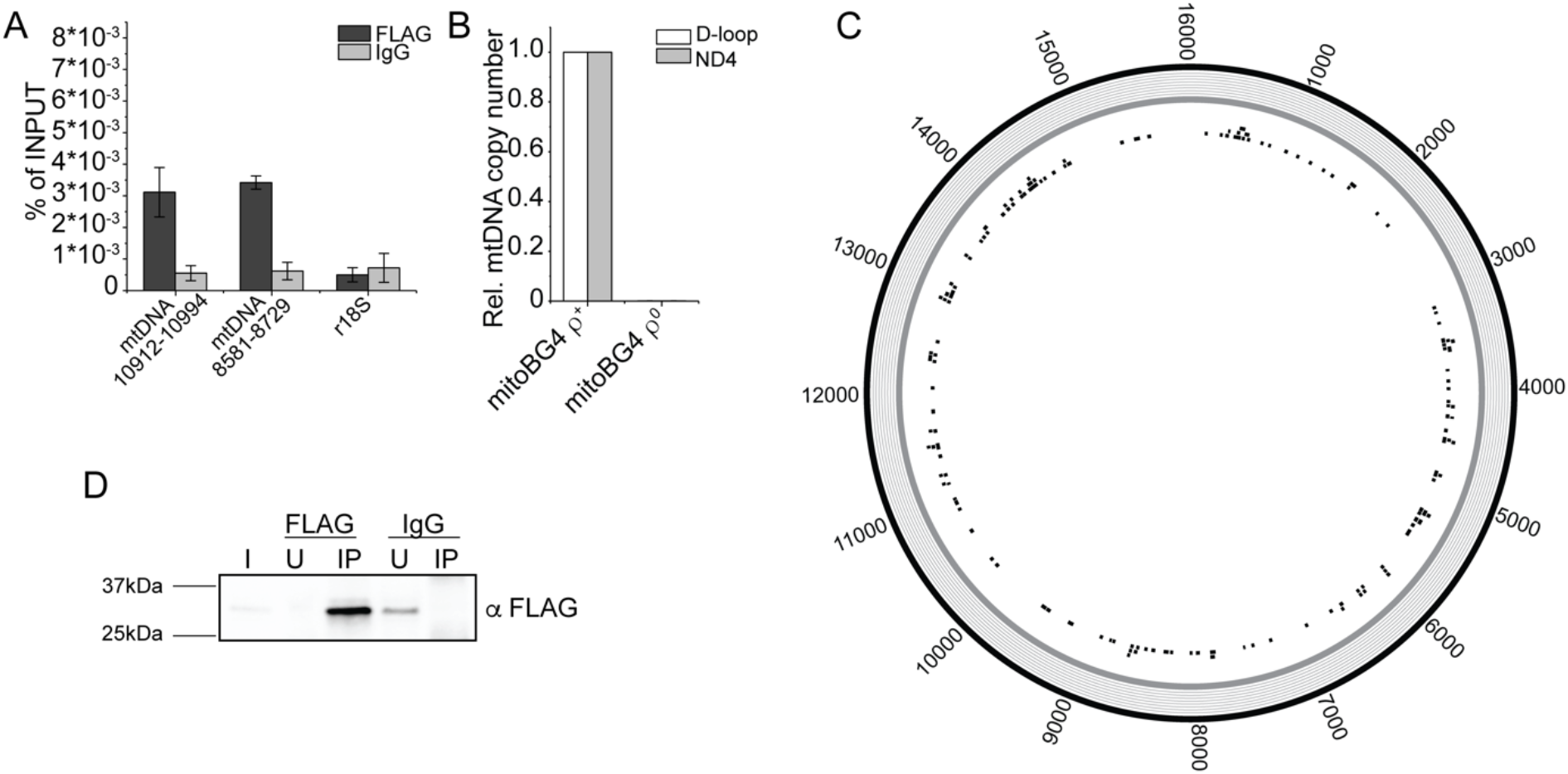
A. ChIP-qPCR upon pull down with FLAG and IgG antibodies in induced mitoBG4 cells. Two regions in the mtDNA and the nDNA region r18s were analyzed. Data are expressed as % of INPUT. Data represent mean ± s.d. of three independent experiments. B. mtDNA copy number of mitoBG4 ρ^+^ and ρ^0^ cells. MtDNA copy number was determined by qPCR from total DNA. Two different regions of the mtDNA, D-loop and ND4, were analyzed. The relative mtDNA copy number was expressed as the ratio between mtDNA regions and nuclear DNA B2M region and normalized to ρ^+^ samples. Data represent mean ± abs error of two independent experiments. C. mtG4-ChIP-seq profile for induced mitoBG4 ρ^0^ cells (lacking the mtDNA) pulled down with FLAG antibody. Grey color indicate no enrichment. The black boxes in the inner circle are predicted G4 sequences (using the G4 hunter algorithm). D. Immunoblot of Input (I), unbound (U) and pull-down (IP) fractions from FLAG and IgG immunoprecipitation of induced mitoBG4 ρ^0^ cells. The blot was probed with the FLAG antibody. FLAG pull down fraction is enriched for mitoBG4.

**Sup.Figure 6:**
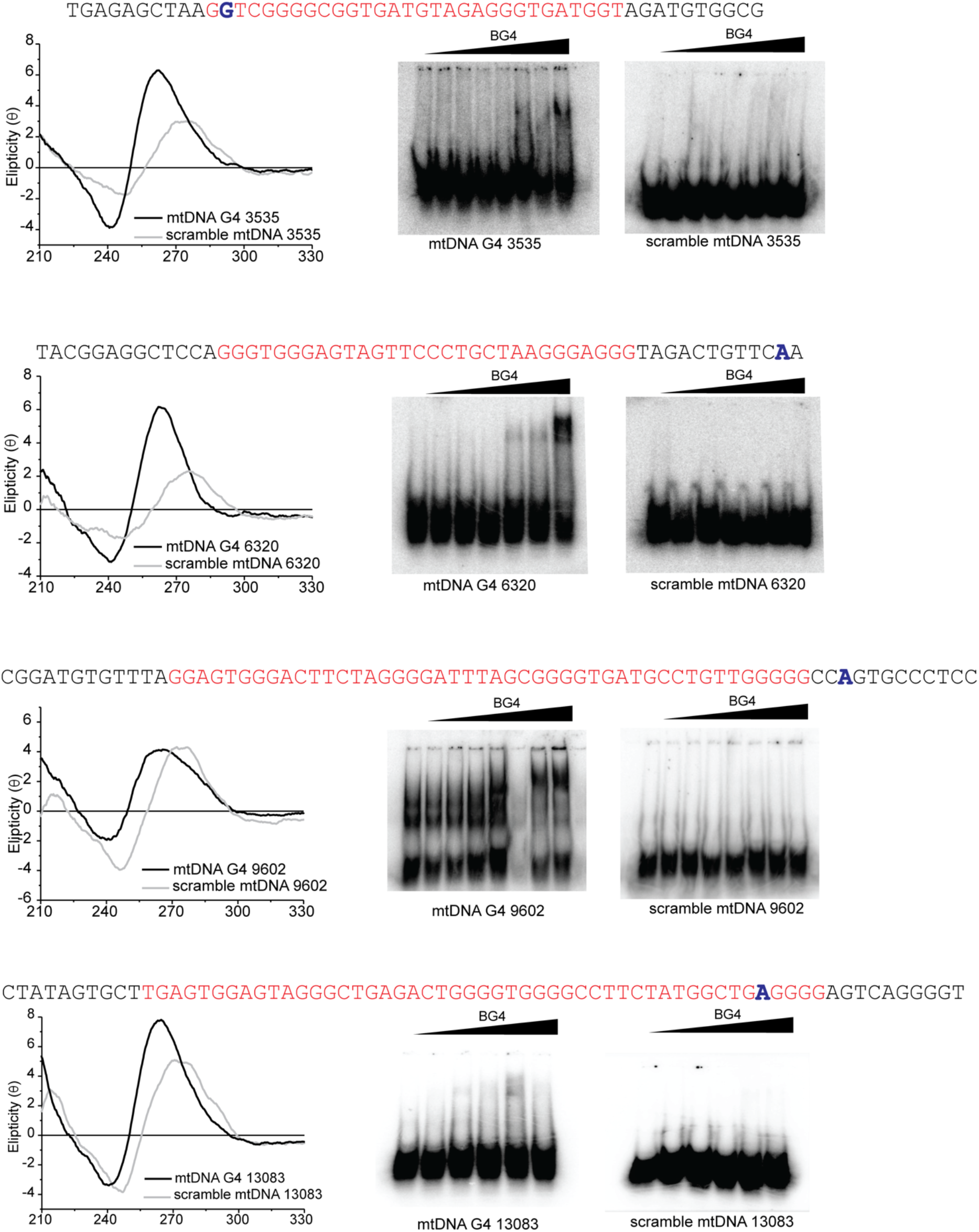
Left: CD spectra of putative G4-forming sequences identified in the mtG4-ChIP-seq protocol. 3 mM of template was folded in 100 mM KCl prior to spectra recording. Scrambled sequences with same guanines content and length were used as control. All PGS analyzed displayed a positive peak at around 264 nm indicative of a G4 structure with parallel topology. Scrambled sequences (grey), present with a spectrum typical of dsDNA. Right: EMSA in the presence of increasing concentrations of recombinant BG4 (BG4) of the oligos used for the CD analysis. DNA concentration was 1 nM. Analyzed sequences are in red, bold letter indicate the peak summit as detected in mtG4-ChIP-seq analysis (see Sup. Table 1).

**Sup.Figure 7:**
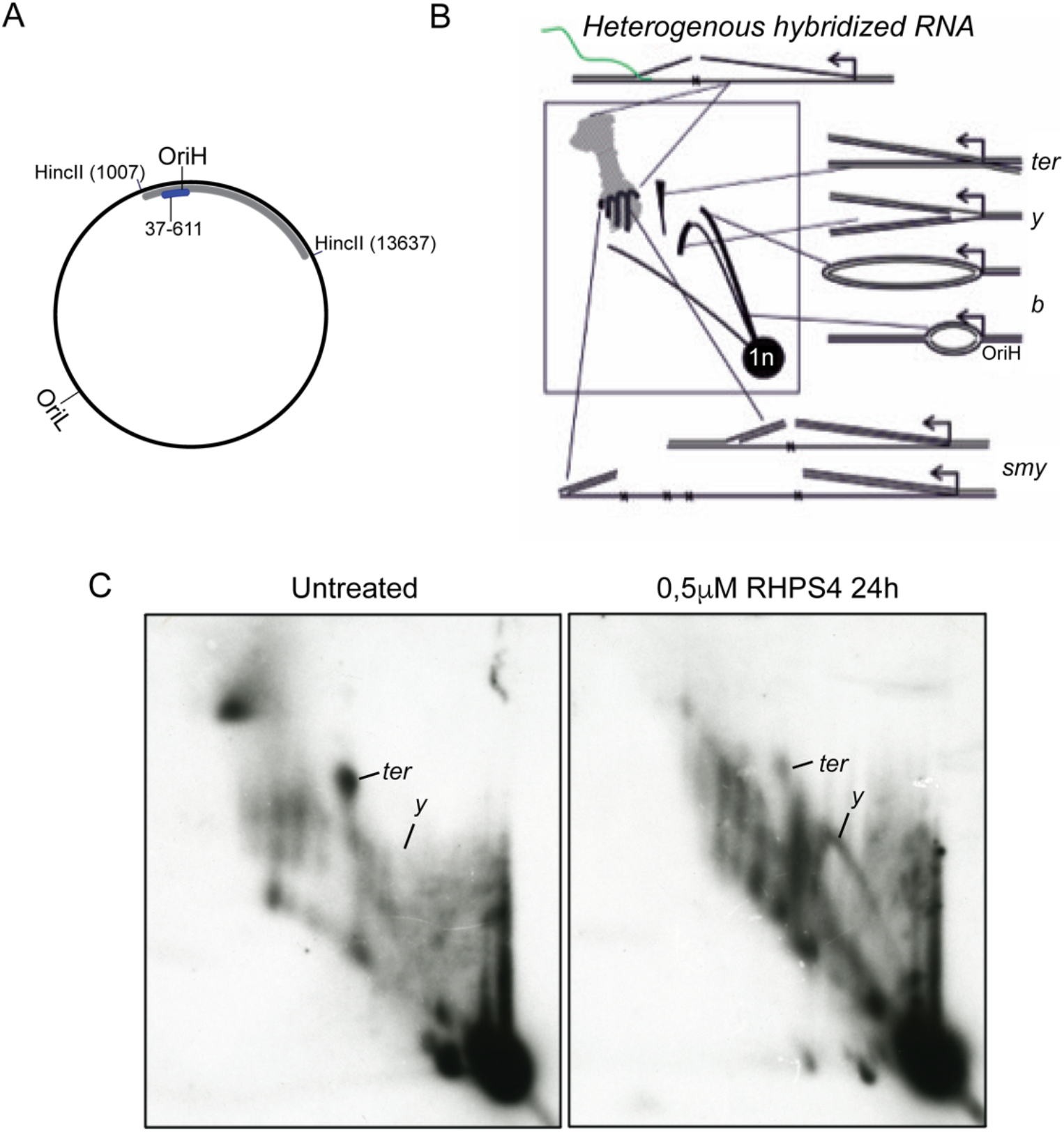
2D-AGE of *Hinc*II digested human mtDNA. A. Schematic illustration of human mtDNA showing the *HincII* restriction sites flanking the non-coding region (NCR). Other *HincII* sites omitted. B. Interpretation of the *Hinc*II NCR 2D-AGE pattern. Linear restriction fragments migrate as a dominant 1n spot. Replication initiating at OH forms replication bubbles, which migrate at the bubble arc (*b*). These convert to replication forks (*y*-forms) and so-called slow moving y-forms (*smy*’s). The latter are generated by incomplete restriction of the displaced H-strand and their size depends on the number of uncut *HincI*I sites (x) they contain. The replication forks meet at the NCR after the replication has completed and form a termination intermediate (ter). C. RHPS4 treatment causes replication stalling, which is visible as enhanced y-arc (*y*) and depletion of the termination intermediates (*ter*).

**Sup.Figure 8:**
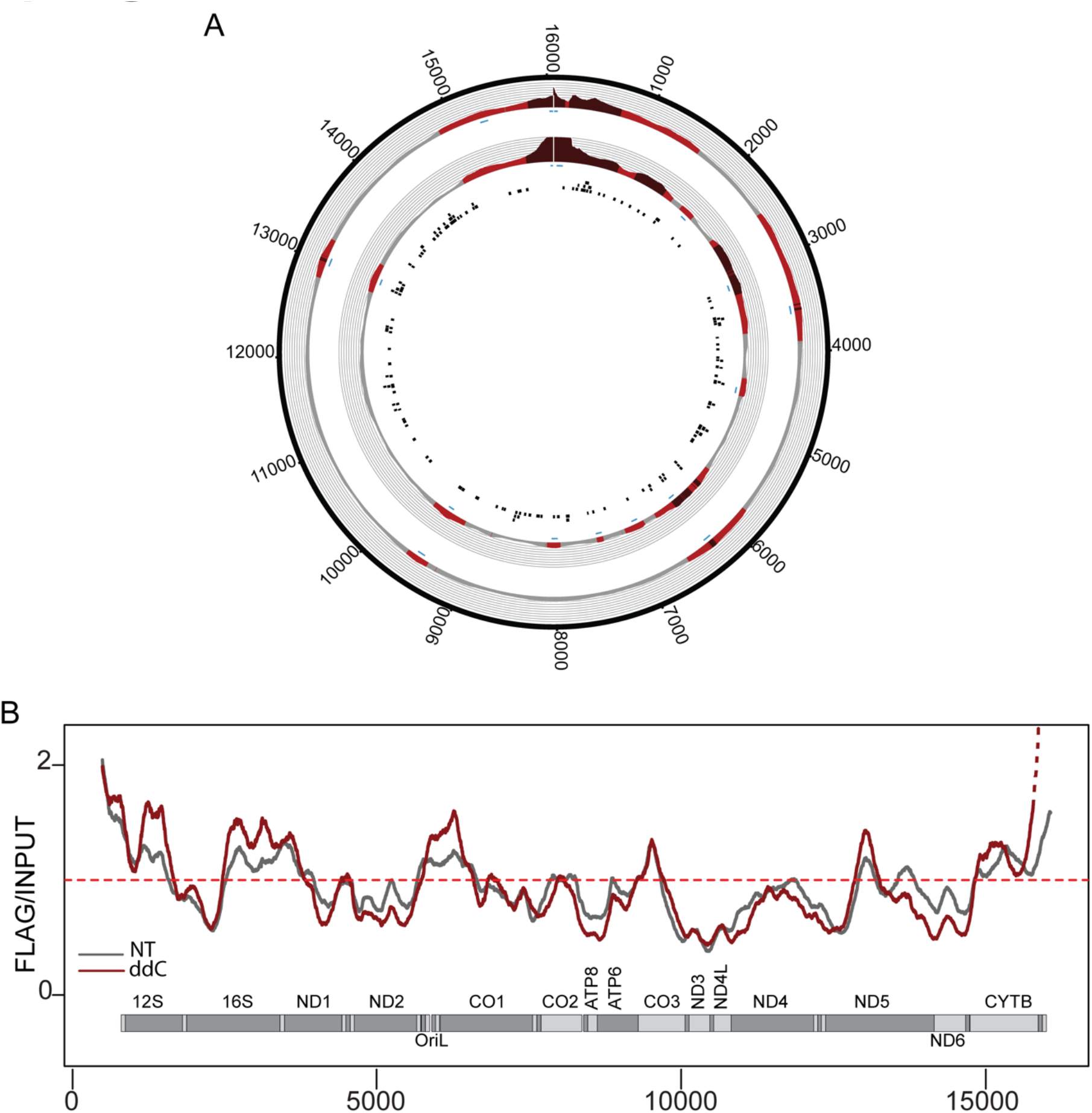
A. mtG4-ChIP-seq profile for induced mitoBG4 cells treated with 200 μM ddC for 3 h before pulling down with FLAG antibody. FLAG signal was normalized to the INPUT sample and expressed as ratio of FLAG vs INPUT. Areas in red indicate enrichment over INPUT samples (gray <1, light red 1 > x > 1,5 and dark red > 1,5). The blue boxes beneath the plots indicates the narrow peaks. The black boxes in the inner circle are predicted G4 sequences (using the G4Hunter algorithm). Two biological replicates are shown. B. Linear plots showing the average FLAG to INPUT signal from the doxycycline induced replicates (in grey) and ddC treated replicates (in red). MtDNA sequence from nt 500 to nt 16 000 is displayed.

**Sup.Table 1:** mtG4-ChIP-seq peak enrichment for doxycycline-induced samples.

| Replicate 1 |  |  | Replicate 2 |  |  | Replicate 3 |  |  | MACS 2<br>PEAK<br>SUMMIT<br>(Rep 1)<br>Flag vs<br>IgG | PGS | PREDICTION |
| --- | --- | --- | --- | --- | --- | --- | --- | --- | --- | --- | --- |
| Peak<br>coordinates<br>(Start – end) |  | Enrichment<br>(Ratio<br>Flag/Input) | Peak<br>coordinates<br>(Start – end) |  | Enrichment<br>(Ratio<br>Flag/Input) | Peak<br>coordinates<br>(Start – end) |  | Enrichment<br>(Ratio<br>Flag/Input) |  |  |  |
| 3072 | 3730 | 1,45 | 2491 | 3989 | 1,40 | 2566 | 3837 | 1,28 | 3528 | MtDNA<br>G4 3535 | G4HUNTER (MITO 27)<sup>a</sup> |
| 6106 | 6681 | 1,20 | 5656 | 6678 | 1,35 | 5839 | 6523 | 1,22 | 6279 | MtDNA<br>G4 6320 | QGRS Mapper<sup>b</sup> |
| 9281 | 9671 | 1,66 | 9128 | 9681 | 1,25 | 9409 | 9584 | 1,12 | 9552 | MtDNA<br>G4 9602 | G4HUNTER (MITO 89)<sup>a</sup> |
| 12937 | 13169 | 1,22 | 12870 | 13852 | 1,30 | 12943 | 13131 | 1,10 | 13034 | MtDNA<br>G4 13083 | G4HUNTER (MITO123)<sup>a</sup> |

<sup>a</sup> Bedrat A, Lacroix L, Mergny JL. Re-evaluation of G-quadruplex propensity with G4Hunter. *Nucleic Acids Res.* 2016;44(4):1746-1759
<sup>b</sup> Kikin O, D'Antonio L, Bagga PS. QGRS Mapper: a web-based server for predicting G-quadruplexes in nucleotide sequences. *Nucleic Acids Res.* 2006 Jul 1;34

**Sup.Table 2:** CD analysis.

| Oligo | Elipcticity $\lambda_{\max}$ (+), $\lambda_{\min}$ (-), nm | Topology |
| --- | --- | --- |
| MtDNA G4 3535 | 262, 241 | G4, Parallel |
| Scramble MtDNA G4 3535 | 275, 248 | Non G4 |
| MtDNA G4 6320 | 263, 241 | G4, Parallel |
| Scramble MtDNA G4 6320 | 275,244 | Non G4 |
| MtDNA G4 9602 | 264, 241 | G4, Parallel |
| Scramble MtDNA G4 9602 | 273, 246 | Non G4 |
| MtDNA G4 13083 | 264, 241 | G4, Parallel |
| Scramble MtDNA G4 13083 | 272, 245 | Non G4 |

**Sup.Table 3:** mtG4-ChIP-seq peak enrichment for ddC and RHPS4 treated samples.

| ddC |  |  |  |  |  | RHPS4 |  |  |  |  |  |
| --- | --- | --- | --- | --- | --- | --- | --- | --- | --- | --- | --- |
| Replicate 1 |  |  | Replicate 2 |  |  | Replicate 1 |  |  | Replicate 2 |  |  |
| Peak coordinates<br>(Start – end) |  | Enrichment<br>(Ratio<br>Flag/Input) | Peak coordinates<br>(Start – end) |  | Enrichment<br>(Ratio<br>Flag/Input) | Peak coordinates<br>(Start – end) |  | Enrichment<br>(Ratio<br>Flag/Input) | Peak coordinates<br>(Start – end) |  | Enrichment<br>(Ratio<br>Flag/Input) |
| 2515 | 3896 | 1,53 | 2455 | 3780 | 1,82 | 2425 | 3789 | 2.46 | 2445 | 3986 | 2.18 |
| 5779 | 6568 | 1,54 | 5672 | 6617 | 1,67 | 5570 | 7168 | 2.01 | 5602 | 7135 | 1.96 |
| 9410 | 9657 | 1,22 | 9261 | 9744 | 1,49 | 9252 | 9837 | 1.58 | 8958 | 9869 | 1.55 |
| 12839 | 13256 | 1,52 | 12866 | 13256 | 1,36 | 12762 | 13307 | 1.37 | 12802 | 13771 | 1.82 |
|  |  |  |  |  |  | 4330 | 4619 | 1.24 | 4350 | 4624 | 1.29 |
|  |  |  |  |  |  | 7182 | 7559 | 1.29 | 7139 | 7534 | 1.38 |
|  |  |  |  |  |  | 7883 | 8244 | 1.41 | 7800 | 8276 | 1.30 |

**Sup. Table 4:**
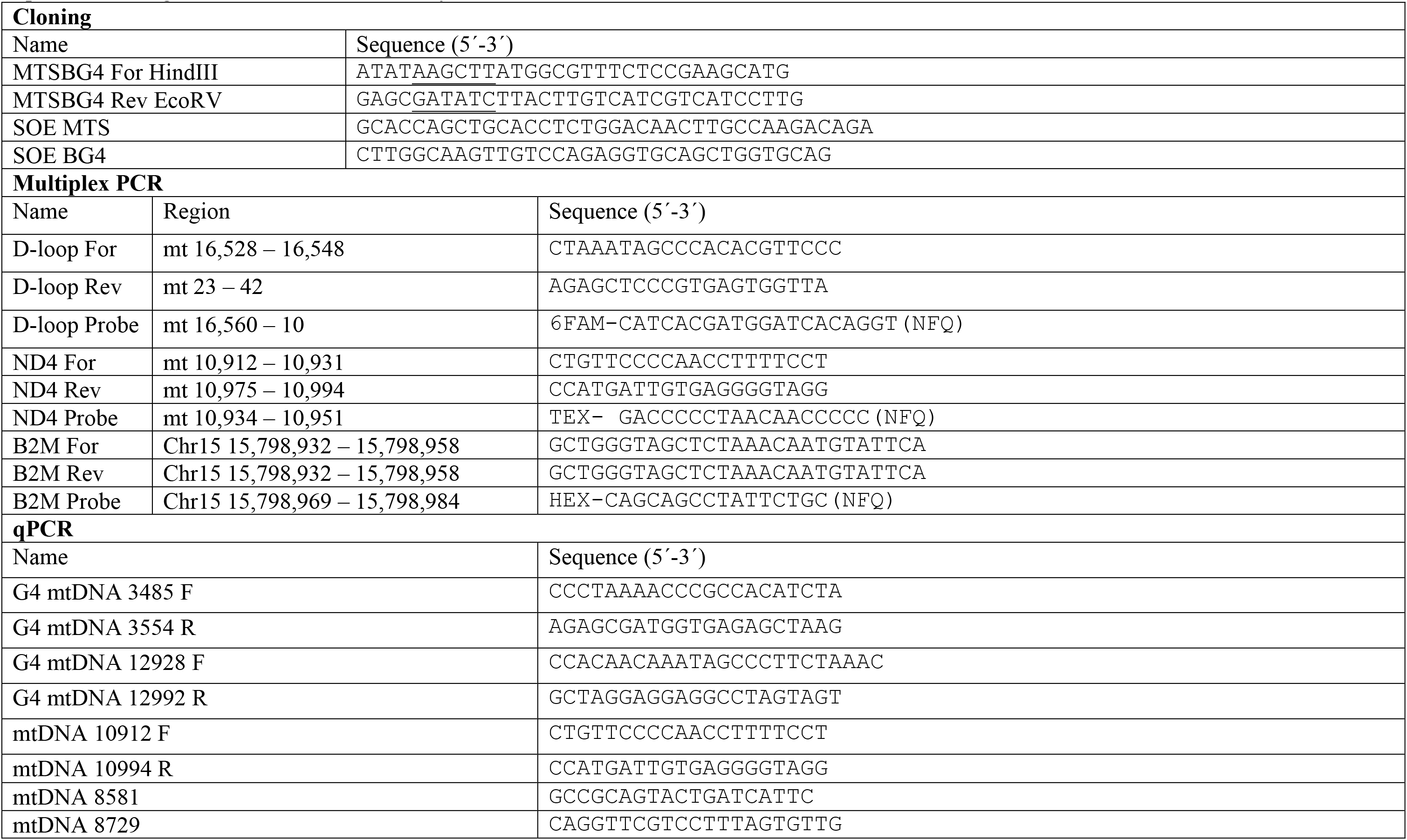
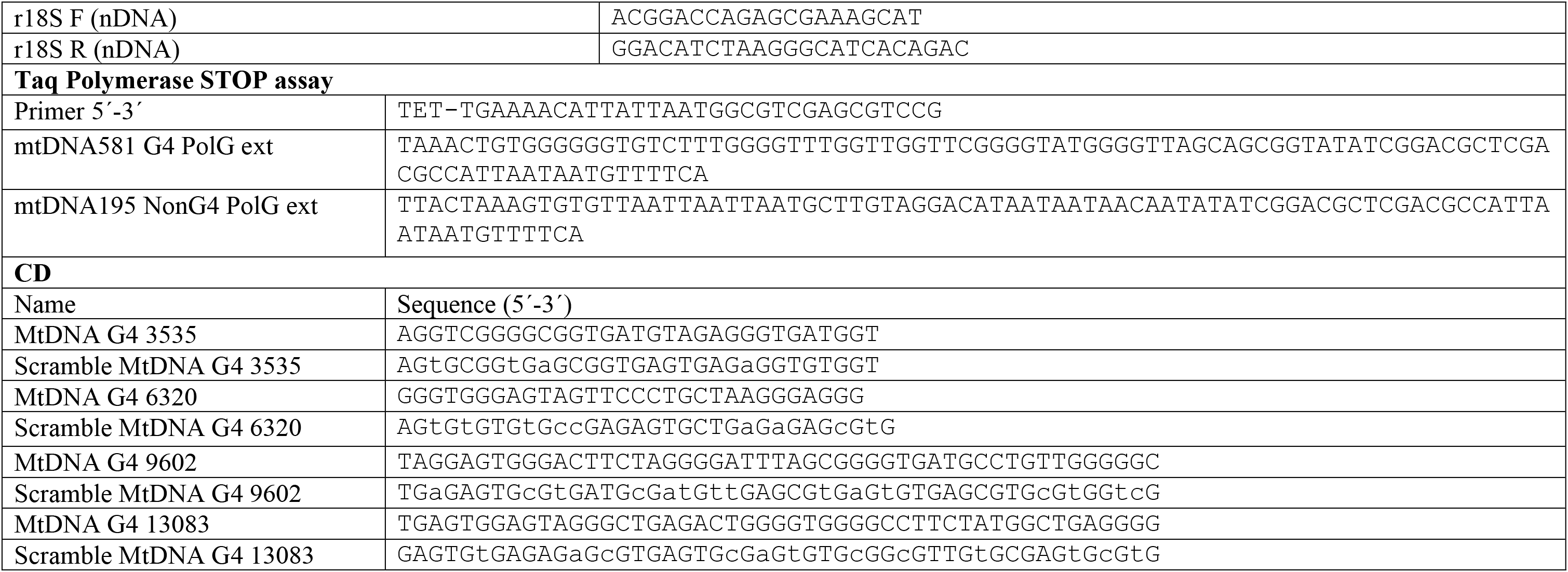
Oligonucleotide used in this study.

**Sup. Table 5:** Antibodies used in this study.

|  |  |  |  |  |  |
| --- | --- | --- | --- | --- | --- |
| *Ab (vs)* | *Company* |  |  | *Cat. num* |  |
| DNA G4 (clone bg4) | Sigma-Aldrich |  |  | MABEE917 |  |
| *Immunoblot (Clean Blot and True Blot antibody used for detection of IP samples)* |  |  |  |  |  |
| *Ab (vs)* | *Company* | *Cat. num* | *dilution* | *II Antibody* | *dilution* |
| Flag M2 | Sigma-Aldrich | F1804 | 1:1000 | Goat anti-Mouse IgG (H+L) HRP (Thermo Scientific) | 1:10000 |
| PolGA | Abcam | Ab128899 | 1:1000 | Clean Blot IP HRP (Thermo Scientific) | 1:1000 |
| B-Tubulin | Rabbit | #2128 | 1:1000 | Goat anti-Rabbit IgG (H+L) HRP (Thermo Scientific) | 1:10000 |
| GAPDH | Cell Signalling | #2118 | 1:1000 | Goat anti-Rabbit IgG (H+L) HRP (Thermo Scientific) | 1:5000 |
| LAMIN A/C | Cell Signalling | #4777 | 1:1000 | Goat anti-Mouse IgG (H+L) HRP (Thermo Scientific) | 1:3000 |
| PrimPol | Customized | ----- | 1:1000 | Clean Blot IP HRP (Thermo Scientific) | 1:1000 |
| B-Actin | Sigma | A5441 | 1:10000 | Mouse TrueBlot ULTRA HRP (Rockland) | 1:2000 |
| C23 MS-3 | Santa Cruz | Sc-8031 | 1:1000 | Mouse TrueBlot ULTRA HRP (Rockland) | 1:2000 |
| Twinkle | Abcam | ab187517 | 1:1000 | Clean Blot IP HRP (Thermo Scientific) | 1:1000 |
| SSBP1 | ProteinTech | 12212-1-AP | 1:1000 | Clean Blot IP HRP (Thermo Scientific) | 1:1000 |
|  |  |  |  | Goat anti-Rabbit IgG (H+L) HRP (Thermo Scientific) | 1:3000 |
| TOM20 | SantaCruz | Sc-11415 | 1:1000 | Clean Blot IP HRP (Thermo Scientific) | 1:1000 |
| TOM20 (F10) | SantaCruz | Sc-17764 | 1:1000 | Goat anti-Mouse IgG (H+L) HRP (Thermo Scientific) | 1:3000 |
| OPA1 | BD Bioscience | 612606 | 1:1000 | Mouse TrueBlot ULTRA HRP (Rockland) | 1:2000 |
|  |  |  |  | Goat anti-Rabbit IgG (H+L) HRP (Thermo Scientific) | 1:3000 |
| TFAM | ProteinTech | 22586-1-AP | 1:1000 | Clean Blot IP HRP (Thermo Scientific) | 1:1000 |
| PCNA | Cell Signalling | #2586 | 1:1000 | Mouse TrueBlot ULTRA HRP (Rockland) | 1:2000 |
| IHC |  |  |  |  |  |
| *Ab (vs)* | *Company* | *Cat. num* | *dilution* | *II Antibody* | *dilution* |
| Flag M2 | Sigma-Aldrich | F1804 | 1:500 | Goat anti Mouse IgG (H+L) –Alexa Fluor594 (Molecular Probes) | 1:1000 |
| TOM20 | SantaCruz | Sc-11415 | 1:200 | Rabbit-488 | 1:500 |
| TFAM | ProteinTech | 22586-1-AP | 1:100 | Rabbit-488 | 1:500 |
| SSBP1 | ProteinTech | 12212-1-AP | 1:100 | Rabbit-488 | 1:500 |

| IP |  |  |  |  |
| --- | --- | --- | --- | --- |
| Antibody | Company | Cat. num | Amount | Host |
| Flag M2 | Sigma-Aldrich | F1804 | 10ug | Mouse |
| Mouse IgG | Santa Cruz | Sc-2025 | 10ug | Mouse |
| PolGA | Invitrogen | PA521314 | 5ug | Rabbit |
| TFAM | Proteintech | 22586-1-AP | 1ug | Rabbit |
| Twinkle | Abcam | Ab187517 | 5ug | Rabbit |
| mtSSB | Proteintech | 12212-1-AP | 1ug | Rabbit |
| PrimPol Neutron | Customized | ----- | 5ug | Rabbit |
| Tom20 | Santa Cruz | SC-11415 | 1ug | Rabbit |
| Rabbit IgG | Santa Cruz | Sc-2027 | 5ug | Rabbit |

